# SketchDNA: A GUI-Enabled Toolkit for Multiscale Modeling of Topological DNA Structures

**DOI:** 10.64898/2026.07.27.741011

**Authors:** Sudhish Gupta, Priyanka Yadav, Himanshu Joshi

## Abstract

Computational modeling tools have enabled detailed exploration of the structural dynamics of nucleic acids at the nanoscale. Despite these developments, a unified platform for creating multiscale models of topological DNA structures, which are of fundamental importance across rapidly converging biological disciplines, is lacking. Here we present a GUI-enabled topological DNA design tool, SketchDNA (`SDNA`), that facilitates a user-friendly interface to create all-atom (AA) and coarse-grained (CG) models of DNA structures with tunable topological parameters. Building on the modular and open-source framework of `SDNA`, the software can be readily integrated with both emerging and existing DNA design platforms, such as oxDNA, MrDNA, *etc*. We demonstrate the utility of `SDNA` computational framework by simulating the conformational dynamics of three representative topological DNA structures, minicircles, catenanes, and Borromean rings. Analyzing AA and CG molecular dynamics (MD) simulations of these topological DNA systems using the AMBER and Martini DNA force fields, respectively, we characterize their equilibrium structural dynamics, fluctuations, and topological properties. While the AA simulations allow us to characterize the topology-dependent structural rearrangements at the nucleotide level, long-timescale CG simulations reveal supercoiling-induced conformational transitions. The multiscale `SDNA` toolkit broadens the applications of MD simulations by enabling *in-situ* characterization of the biophysical properties of topological DNA nanostructures. The GUI version of `SDNA` is available at https://sdna.biotech.iith.ac.in without registration, while the source code is accessible at GitHub

## Introduction

DNA is arguably the most studied biomolecular structure of the last century [1]. From a single nucleotide (0.34 nm, rise per base-pair) to the complete genome (∼ 2 meter, typical contour length of DNA in a biological cell) the structural organization of DNA spans an extraordinary length scale. Although the canonical B-form of DNA represents the thermodynamically stable ground-state conformation of DNA, its geometry continuously fluctuates around this equilibrium structure due to torsional and tensile stresses during various cellular processes. These deformations often lead to over-twisted or under-twisted structures, commonly known as supercoiled DNA [2]. DNA molecules inherently experience topological tension arising from torsional and spatial constraints generated during DNA replication, transcription, recombination, chromosomal segregation, repair, and other cellular processes [3**?**]. Topological constraints combined with thermal fluctuations lead to torsional stress that induces supercoiling of the strands and eventually drives DNA to adopt diverse structural conformations [4, 5]. Representative biological examples include plasmids, chromatin loops, nucleosomes, and mitochondrial genomes, whose topology is dynamically regulated by proteins such as topoisomerases [6], histones, and nucleoid-associated proteins [7] (e.g. H-NS) to preserve genome organization and function [8].

Inspired by the precise and programmable structural organization of naturally occurring DNA systems, advances in DNA nanotechnology have led to the construction of custom-shaped nanostructures that expand beyond their biological functions into the material world [9, 10, 11**?**]. Explorations in DNA nanotechnology have also led to the construction of topological DNA architectures such as minicircles [12], catenanes [13, 14], Möbius strip [15], rotaxanes [16], Borromean rings [17] and other chiral DNA systems [18]. These topological DNA nanostructures have been proposed for their applications in gene editing [19], nanosensing [20], drug delivery and programmable molecular systems [21] etc.

DNA minicircles, catenanes, and Borromean rings present archetypal topological DNA systems that can provide fundamental insights into genome regulation, topoisomerase mechanism, and rational design of interlocked DNA nanoarchitectures. For example, DNA minicircles with tunable superhelical density have been very useful for studying DNA plasmids [12]. Similarly, DNA catenanes can mimic interlocked structures that occur transiently in living cells during DNA replication and homologous recombination and are resolved by topoisomerases to ensure faithful chromosome segregation [6]. The programmable self-assembly and mechanical interlocking of DNA catenane have also made them attractive building blocks for DNA nanotechnology, where they have been incorporated into functional molecular devices, such as molecular switches, nanomachines, responsive materials, etc [14, 20]. Borromean rings [22] represent a unique and interesting higher-order topology in which three rings are collectively interlocked while no two rings are directly linked. Their successful realization through DNA self-assembly marked an important milestone in structural DNA nanotechnology, demonstrating the ability of DNA to form sophisticated mechanically interlocked architectures [17].

Advancements in experimental techniques, especially the single-molecule imaging methods, have enhanced our understanding of topological DNA nanostructures[23, 24, 25]. At the same time, all-atom (AA) molecular dynamics (MD) simulations in the exascale computing era have been able to reveal the critical biophysical properties of nucleic acids at atomic resolution, complementing experimental observations by providing molecular insights that are difficult to characterize in experiments [26, 27, 28, 29, 30, 31, 32**?**]. Coarse-grained (CG) models, such as Martini [33] and oxDNA [34], overcome the limitations of length and timescale associated with AA simulation by reducing molecular resolution, and thereby allowing simulations of large DNA assemblies for longer times. Integrating AA and CG simulations provides a powerful framework allowing accurate characterization of molecular interactions while capturing long-timescale conformational dynamics.

Setting up multiscale MD simulations requires reliable tools for the consistent and flexible generation of molecular configurations and topologies. Apart from the quality of the force fields, the accuracy of the molecular simulations often depends on the initial starting structure. Several softwares have been developed for constructing and analyzing nucleic acid structures. General-purpose builders such as Nucleic Acid Builder (NAB) [35], 3DNA [36], and the recently developed MDNA framework [37] facilitate the generation of canonical DNA structures and provide extensive support for sequence-dependent modeling and structural analysis. In parallel, advances in DNA nanotechnology have led to the development of specialized design platforms, including caDNAno [38], Tiamat [39], CanDo [40], vHelix [41], DAEDALUS[42], Adenita [43], MagicDNA [44], and DNA forge [45] which enable the construction of increasingly complex DNA origami and wireframe nanostructures. Complementary tools such as MrDNA [46] provide multiresolution modeling for the design and structural relaxation of DNA origami assemblies, while coarse-grained frameworks such as oxDNA [34, 47] facilitate the simulation and analysis of large DNA systems.

However, none of the frameworks developed so far provide a straightforward workflow for generating multiscale models of custom-shaped topological DNA architectures, hindering their computational exploration. Most existing modeling tools are designed for linear duplex DNA and require extensive manual manipulation or scripting to generate non-linear DNA topologies with tunable superhelical densities. Preparing these systems for all-atom and coarse-grained simulations is often time-consuming, error-prone, and requires expertise in multiple software packages [48, 49].

Here, we present SketchDNA(`SDNA`), a GUI-based open-source toolkit built on the MDNA framework [37] for the automated generation of simulation-ready DNA models in both AA and Martini CG representations. `SDNA` enables the construction of a broad range of customizable DNA architectures with user-defined geometrical and topological parameters, including the option to tune the superhelical density and sequence. Figure 1 illustrates the complete workflow of the `SDNA` toolkit, and Supplementary Movie 1 provides a video tutorial demonstrating the step-by-step usage of the tool. To exemplify the capabilities of `SDNA`, we generated three representative DNA topologies and performed multiscale simulations of DNA minicircles with varying superhelical densities, DNA catenanes, and DNA Borromean rings. Following the introduction, we describe the design and implementation of the `SDNA` framework, including the construction of topological DNA architectures and the development of simulation protocols. Finally, we present the results from multiscale MD simulations of three representative topological DNA nanostructures and discuss the capabilities, current limitations, and potential future applications of `SDNA`.

**Figure 1.**
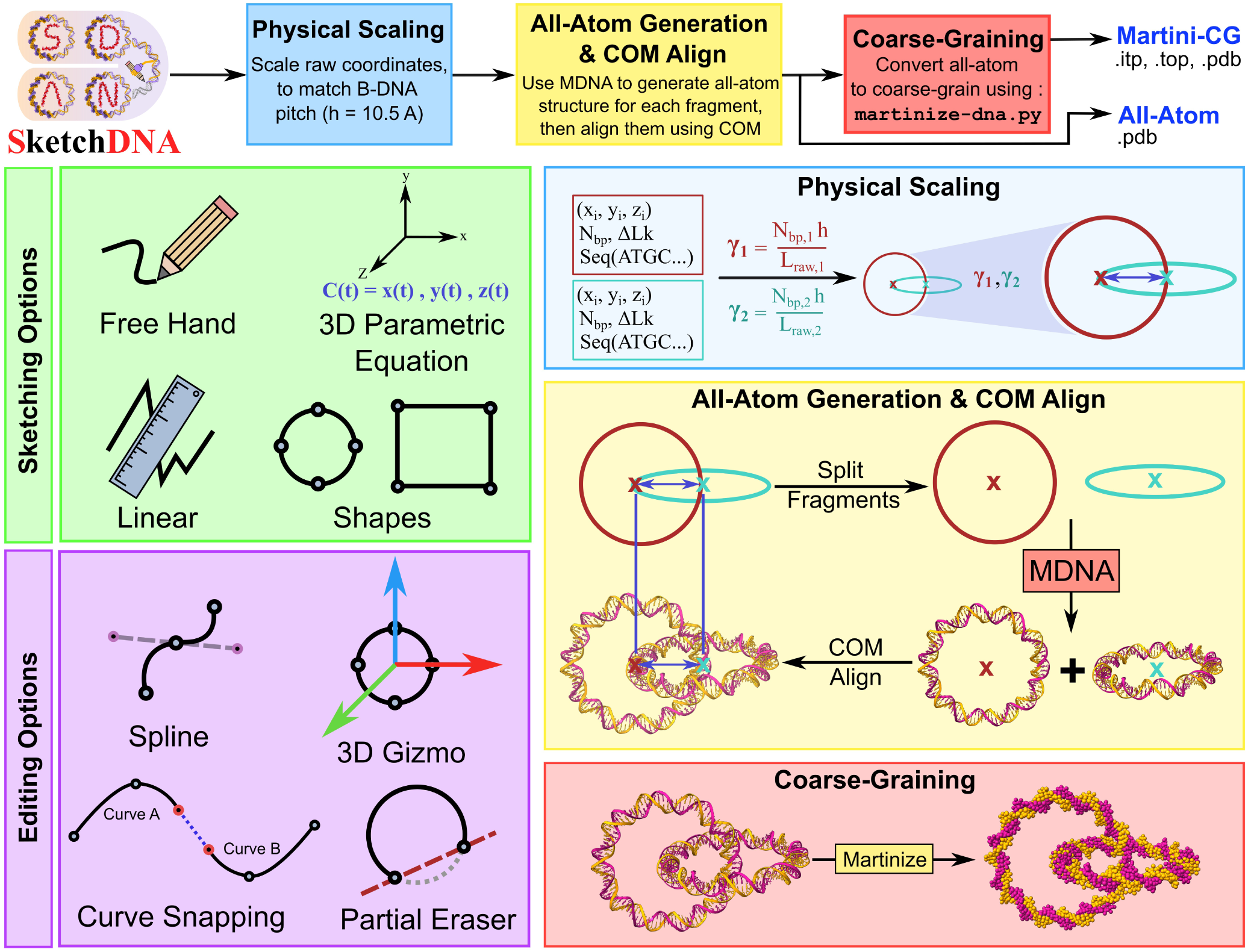
Overview of the SDNA pipeline to obtain multiscale models of topological DNA systems. The user can utilize a variety of input options and GUI-enabled editing features to sketch their desired DNA topology with tunable supercoiling densities. The sketched design is transmitted as 3D Cartesian coordinates, which are further scaled to ensure the design can accommodate the requested number of base-pairs. MDNA [37] is then used to process these scaled coordinates to generate the AA structure for each individual fragment. Further, the AA model is coarse-grained using the Martini protocol. Structure can be visualized and downloaded from `SDNA` for further simulations using MD simulation suit.

## Materials and Methods

### SDNA Framework Architecture

The `SDNA` platform is an integrated multi-scale computational pipeline designed to facilitate the intuitive three-dimensional (3D) models of complex ds-DNA nanostructures. The framework utilizes a coupled architecture consisting of a `React.js` based web interface to sketch the designs, supported by a `python` based backend to convert the designs into all-atom ds-DNA models using MDNA and further coarse-grain them into Martini beads.

#### User Interface and Geometric Design

The web-interface provides the user with a localized workspace for the spatial orchestration of complex DNA architectures. The interface supports a diverse array of geometric primitives and free-form sketching modalities:

- **Sketching Modalities:** Users may instantiate DNA paths through free-hand drawing, linear segments, or cubic spline-based curved segments.
- **Parametric Architectures:** Input modalities also support regular polygons (e.g., squares, triangles, *n*-sided polygons) with adjustable border radii, which prevent the formation of sharp kinks in regular geometric shapes. `SDNA` also supports the design of complex 3D paths defined by user-specified parametric equations.
- **Spatial Manipulation:** The tool also facilitates spatial manipulation by enabling drag and drop 3D translation of fragments.
- **Combining Separate DNA Fragments** Users can combine two separate DNA fragments using an automated “snapping” algorithm to join proximal endpoints, ensuring topological continuity in multi-fragment constructs.
- **Visualization** Users can also view the generated structures in real-time within the `SDNA` interface. The toolkit uses `3Dmol.js` to provide separate dedicated 3D viewer panels for both AA and CG resolution of the system, with several options for molecule representation and coloring scheme. This allows the user to directly compare their design with the generated structures without switching between `SDNA` and other molecular-visualization tools.

#### All-Atom Structure Generation using MDNA

Upon finalization of the geometric design, the spatial coordinates and global parameters for each fragment – including base pair count (*N*_bp_), linking number (ΔLk) which controls the superhelical density, and the desired nucleotide sequence–are transmitted to the backend for generating the structures.

- **Physical Scaling:** To ensure the structural model accurately reflects the physical dimensions of B-form DNA, the input coordinates for each curve fragment is scaled isotropically. We define a separate scale factor *γ_i_* for each curve fragment *i* that maps the discrete arc length of the control points (*L_raw_*|*_i_*) to the theoretical contour length of the DNA duplex.

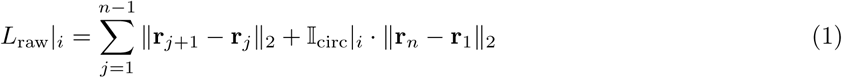 where **r***_j_* ∈ ℝ^3^ are the *n* control points for the curve fragment *i* and ‖_circ_|*_i_* ∈ {0, 1} is an indicator if the fragment is closed/circular. Given a target count of *N*_bp_|*_i_* base pairs for the curve fragment *i* and a canonical B-DNA helical rise *h* ≈ 3.4 Å, the dimensionless scaling factor *γ_i_* is defined as:

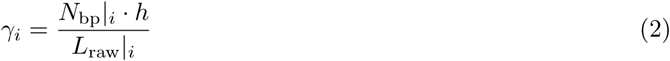
- **All-Atom Model Generation:** High-resolution all-atom structures are generated for each individual fragment utilizing the MDNA[37] framework. The transformed coordinates 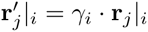 allows MDNA to interpolate points for inserting the requested number of base-pairs (*N*_bp_) along the prescribed trajectory, enabling the user to easily reuse the same geometric design, for any desired size of the system.
- **Spatial Integrity:** For multi-fragment constructs, `SDNA` processes each fragment separately to generate the respective individual structures. To preserve the integrity of the user-defined architecture, the Center of Mass (COM) of each generated atomic fragment *i* is then translated to align precisely with the COM derived from the *γ_i_* − scaled design coordinates of the fragment *i*.

#### Coarse-Grained Mapping

To study longer time-scale evolution of the designed nanostructures, SDNA provides a coarse-grained version of the full atomistic design using an automated coarse-graining pipeline:

- **Martini Conversion:** The consolidated AA structure is processed *via* a modified version of `martinize-dna.py`[50] script. The modification enables coarse-graining of both open / closed DNA structures, by refactoring the backbone bead placement module of the original script.
- **Topology Generation:** The script generates **Martini2** coarse-grained DNA models [51], accompanied by a global topology (`.top`) file and a single Include Topology (`.itp`) file for all the DNA chains present in the design.

#### Output Files

`SDNA` stores AA and CG structures as separate `.pdb` files. Apart from this the `.top` and `.itp` files required for Martini CG simulations are also provided. While exporting, `SDNA` bundles all the files into a `single .zip` file that the user can easily download to their system.

### Construction protocol for selected DNA Topological Structures

To demonstrate the ability to produce high-fidelity multi-resolution models, we employed `SDNA` to generate a diverse set of topological DNA structures that span multiple levels of complexity (Figure 2). All these designs were created using various input options or combinations of the same.

**Figure 2.**
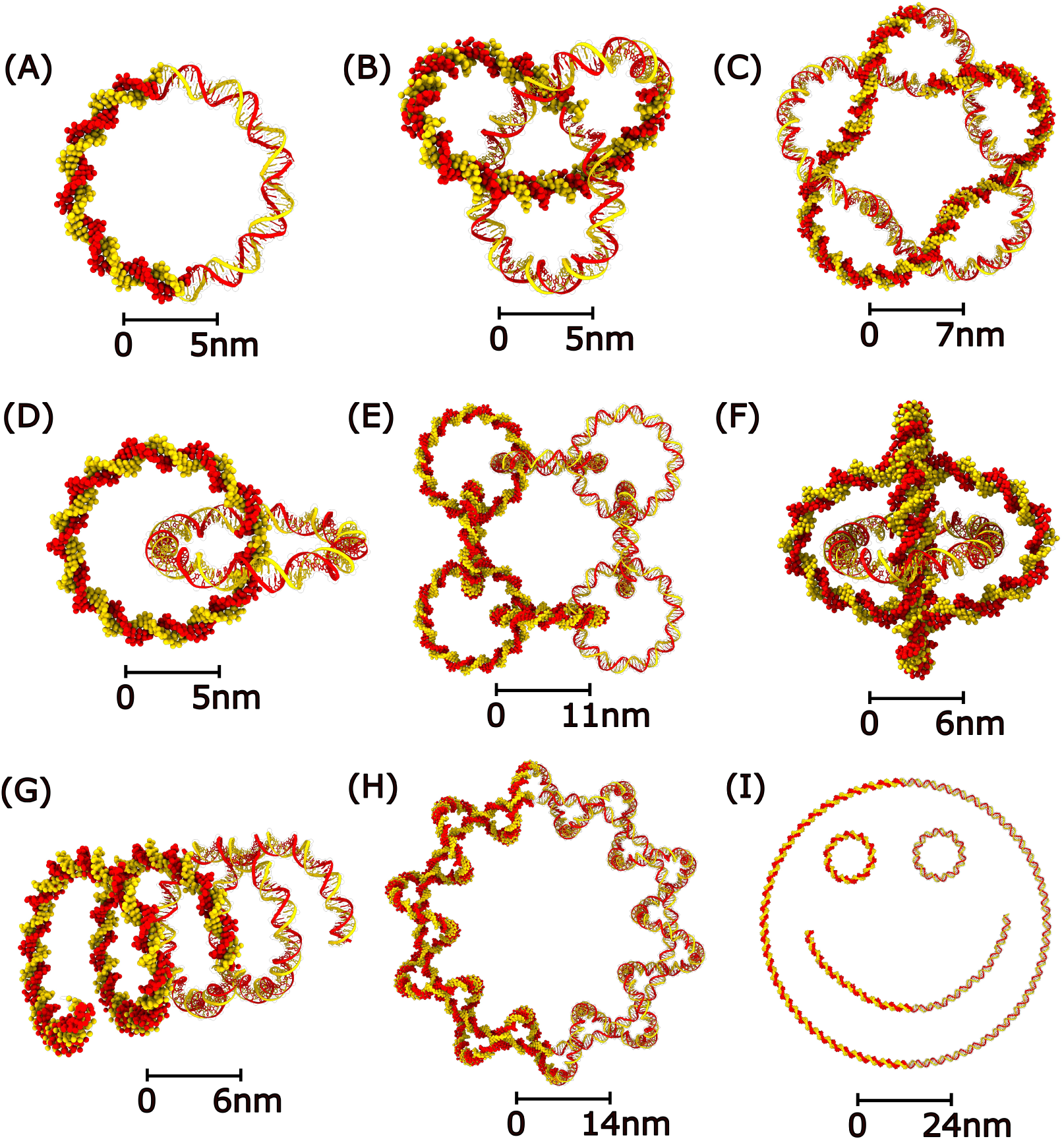
Representative DNA nanostructures generated using SDNA. Illustrating the versatility of the multiscale `SDNA` platform for constructing a wide range of topologically complex DNA architectures. The structures include (A) Minicircle, (B) Trefoil knot, (C) Pentafoil knot, (D) Homocatenane, (E) Polycatenane in a closed-loop, (F) Borromean ring, (G) Solenoidal (helical) DNA structure, (H) Minicircle threaded through a toroidal spring, and (I) Smiley-face DNA nanostructure. For each panel, the AA model is shown alongside its corresponding Martini CG representation purely to illustrate the multiscale versatility of `SDNA`. Approximate scale bars are added to each panel to show the respective length scale.

Below, we provide the building protocol of the three selected topological DNA structures simulated to illustrate the utility of SDNA. All constructs were generated using parametric representations of closed curves in ℝ^3^. Unless otherwise stated, *c* ∈ ℝ denotes a constant offset along one axis, and *R, R*_1_*, R*_2_ denote radii. The parameter *t* belongs to [0, 2*π*]. Although the `SDNA` pipeline supports the generation of DNA structures with custom sequences, we used a poly-AT sequence across all the simulated systems for simplicity.

#### DNA Minicircle

A parametric equation of DNA minicircle (Figure 2 A) is defined as:

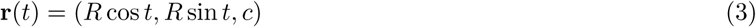

We created 3 separate constructs, each of *N*_bp_ = 100 base pairs, but with ΔLk = 0 for no supercoiling, ΔLk = +1 for positive supercoiling and ΔLk = −1 for negative supercoiling. The radius is set to *R* = 1.0.

#### DNA Catenanes

DNA catenanes (Figure 2 D) are constructed as interlinked closed curves with orthogonal orientations.

#### Heterocatenane

Two minicircles of unequal size are defined as:

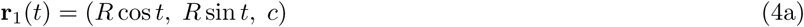

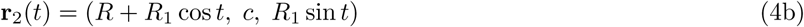

The first and second rings consist of *N*_bp_ = 200 and 100 base pairs, respectively, both with ΔLk = 0. The radii are set to *R* = 2.0 and *R*_1_ = 1.0.

#### Homocatenane

As a special case of Hetrocatenanes, two minicircles of identical size, termed as Homocatenanes are defined as:

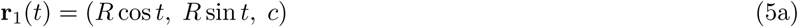

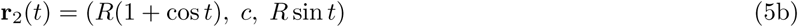

Each ring consists of *N*_bp_ = 100 base pairs with ΔLk = 0. The radius of each ring is set to *R* = 1.0.

#### Borromean rings

Three mutually interlocked elliptical rings in a Borromean setup (Figure 2 F) are constructed as:

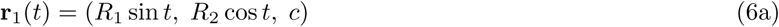

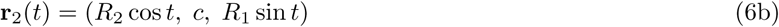

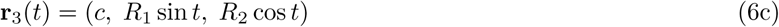

All 3 rings consist of 120 base pairs, with ΔLk = 0. The radii are set to *R*_1_ = 0.75 and *R*_2_ = 1.25.

### Protocols for explicit solvent multiscale MD simulations

The AA models obtained from `SDNA` can be simulated using any MD simulation engines like, AMBER [52], GROMACS [53], or NAMD [54] *etc.* using LEaP, gmx or psfgen tools respectively while the CG Martini models are readily available for running in GROMACS. AA and CG MD simulations were performed for selected DNA topologies in this study, namely minicircles, catenanes, and Borromean rings. The initial structures generated using `SDNA` were subsequently prepared for simulation using AMBER24 [52] and GROMACS [53] for AA and CG models, respectively. In both cases, DNA systems were solvated in explicit water, neutralized with Na^+^ ions, and maintained at 150 mM salt concentration of NaCl. AMBER force fields [55] updates were used to describe the AA representations in AMBER, whereas the CG systems were modeled using the Martini2 force field and soft elastic network [33, 51]. A detailed description of system preparation, force fields, protocols for equilibration and production simulations is provided in Supplementary Information, section S1, while Supplementary Information, section S2, gives the details of various analyses and definitions used in the article. Table 1 describes the details of the simulated systems.

**Table 1.** Details of the simulated topological DNA systems using the AA and Martini CG models obtained from `SDNA`. Here, ΔLk represents the change in the linking number, while *σ* denotes the superhelical density as defined in Equation (7) and (8). Values in parentheses indicate the number of DNA atoms in the AA models and DNA beads in the CG models, respectively. For the sake of simplicity and homogeneity, we used only the **poly-AT** sequence across all the AA and CG models.

| Topology | Length (bp) | $\Delta\text{Lk}$ | $\sigma$ | AA atoms | CG beads |
| --- | --- | --- | --- | --- | --- |
| Minicircle | 100 | 0 | 0.0 | 193,986 (6,400) | 31,844 (1,300) |
|  | 100 | +1 | +0.1 | 194,802 (6,400) | 31,598 (1,300) |
|  | 100 | -1 | -0.1 | 195,354 (6,400) | 53,181 (1,300) |
| Homocatenane | 100 + 100 | – | – | 528,678 (12,800) | 85,770 (2,600) |
| Heterocatenane | 200 + 100 | – | – | 1,344,213 (19,200) | 245,071 (3,900) |
| Borromean ring | 120 + 120 + 120 | – | – | 913,281 (23,040) | 124,495 (4,680) |

## Results and Discussions

### Multiscale simulations reveal superhelical density-dependent structural plasticity of DNA minicircles

DNA minicircles provide an excellent platform for studying plasmid DNA and exploring their potential applications in gene therapy [12, 29]. In a circular DNA, the superhelical stress can modify the equilibrium twist (Tw) which is compensated by the introduction of writhe (Wr) in the polymer, to keep linking number (Lk) constant [56]. Formally, these 3 topological descriptors of a DNA minicircle are mathematically related as follows :

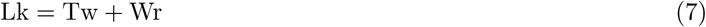

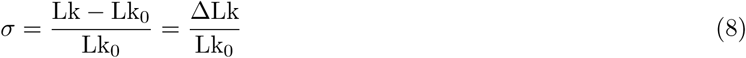

where *σ* denotes the supercoiling density, ΔLk = Lk − Lk_0_ is the change in the linking number relative to relaxed DNA, and Lk_0_ is the linking number of the relaxed DNA molecule. Using `SDNA`, we generated DNA minicircles at three pre-defined superhelical densities, relaxed (*σ* = 0), positive (*σ* = +0.1), and negative (*σ* = −0.1) supercoiling, Table 1. Supplementary movie SM2 illustrates the conformational evolution of all three DNA minicircles during the 0.5*µ*s AA and 10*µ*s Martini CG simulations.

Figure 3 A–C shows the conformation of the DNA minicircles at the beginning and end of the 0.5*µ*s AA (top row) and 10*µ*s of CG (bottom row) simulations, respectively. The relaxed minicircle mostly maintains its circular geometry in AA simulations, exhibiting slightly higher thermal fluctuations in the CG simulations (Figure 3 A). In contrast, both positively and negatively supercoiled minicircles undergo pronounced structural rearrangements as characterized by root mean square deviation (RMSD) and radius of gyration (R*_g_*), (Figure 3 B,C and Supplementary Figure S3 A–D). The positively supercoiled minicircle develops localized kinks, whereas the negatively supercoiled minicircle exhibits transient local strand separation accompanied by disruption of Watson–Crick base pairing (Figure 3 B, C, top). The longer CG simulations sample a broader conformational landscape, resulting in stable right-handed and left-handed writhed conformations for the positively and negatively supercoiled minicircles, respectively (Figure 3 B,C, bottom).

**Figure 3.**
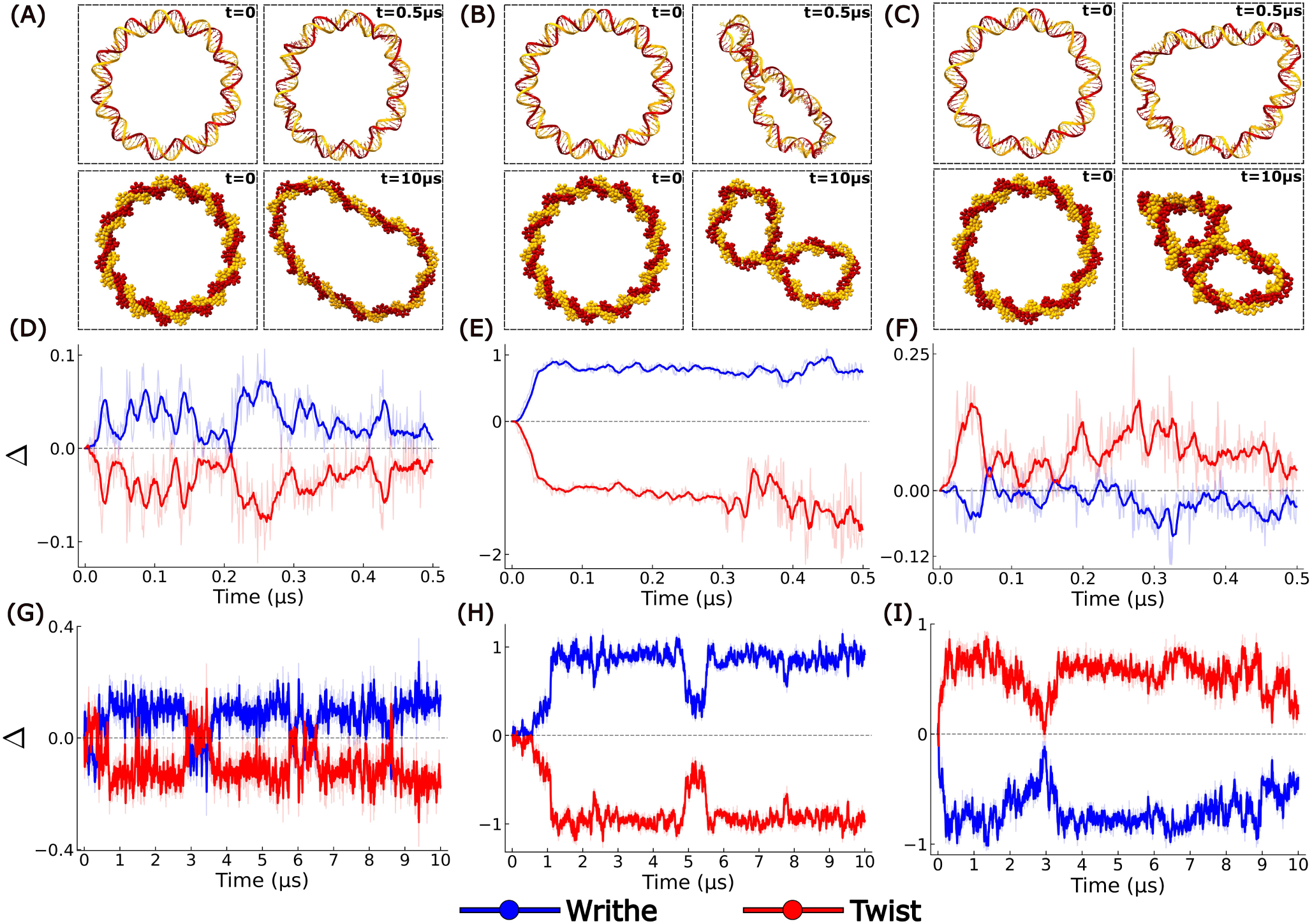
Structural and topological evolution of DNA minicircles with varying superhelical parameters. (A–C) Representative snapshots of DNA minicircles with superhelical densities *σ* = 0 (left), *σ* = +0.1 (middle), and *σ* = −0.1 (right) at the beginning and end of AA (top) and CG (bottom) simulations. Water and ions are not shown for the sake of clarity. Time evolution of change in twist (Δ*Tw*) and writhe (Δ*Wr*) as a function of simulation time in (D–F) AA models and (G–I) CG models for *σ* = 0, *σ* = +0.1, and *σ* = −0.1, respectively.

The analysis of Watson–Crick base pairing revealed that the relaxed minicircle maintained a higher average number of hydrogen bonds (H-bonds) than the positively and negatively supercoiled minicircles, (Supplementary Figure S4). This indicates duplex stability in the relaxed state. The reduced number of H-bonds observed in the supercoiled systems is consistent with transient base-pair disruption, which has previously been associated with formation of localized kinks and denaturation bubbles that relieve bending and torsional stress in minicircles [57, 58, 59, 60]. Figure 3 D-I illustrates the evolution of superhelical stress in minicircles characterized by the change in the topological descriptors, twist (ΔTw) and writhe (ΔWr) during simulations. WrLINE [61] was used to calculate writhe for AA simulations, whereas for CG simulations, an in-house modified version of WrLINE was used (Supplementary Information section S2(C)).

For the relaxed minicircle both AA and CG simulations exhibit only small fluctuations in twist and writhe throughout the simulations, indicating minimal redistribution between the two topological components and preservation of the relaxed topology (Figure 3 D, G). The nature of supercoiling determines the chirality of writhing. Positively supercoiled minicircle (*σ* = +0.1) rapidly converts excess twist into positive writhe. In AA simulation, writhe increases within the first ∼ 100 ns and subsequently remains nearly constant, accompanied by a corresponding decrease in twist (Figure 3 E). CG simulation reproduces a similar overall behavior over a longer timescale, with writhe gradually increasing and stabilizing despite broader conformational sampling (Figure 3 H). In contrast, negatively supercoiled minicircle (*σ* = −0.1) shows persistent fluctuations in both twist and writhe throughout the AA and CG simulations. Unlike the positively supercoiled system, AA simulation does not result in formation of stable negative writhing, while the transient negative writhe observed in the CG simulation gradually decreases towards the end of the simulation (Figure3 I). This redistribution of the conserved linking number (Lk) between twist (Tw) and writhe (Wr) is consistent with previous studies showing that the underwound and overwound supercoiled DNA relieves torsional stress through continuous twist-to-writhe interconversion [4, 60, 62, 63, 64].

The time–averaged distance matrix ⟨*D*⟩ between the COM of the nucleotide base-pairs (Supplementary Figure S5) shows that the relaxed minicircle preserves a nearly symmetric distance pattern at both resolutions, whereas the positively supercoiled minicircle exhibits distinct off-diagonal contacts arising from writhed conformations, consistent with the structural transitions observed in the simulation. AA simulations reveal no significant off-diagonal contacts for negatively supercoiled minicircle. Whereas in longer CG simulations, both the positive and negative superhelical DNA show moderate to high off-diagonal proximity patterns consistent with the writhing observed during the simulations.

The ion density distribution (Supplementary Figure S6,S7) reveals preferential localization of counterions along the DNA backbone across all three topological states. The CG simulation reproduces the overall ion-condensation behaviour but exhibits a more diffuse spatial distribution than the atomistic model. The diffuse ion distribution around DNA in CG simulations could be a manifestation of larger Martini beads that prevent ions from penetrating DNA grooves as previously reported by Naskar *et al.* [65].

Overall, these results show that the atomistic simulations capture the local structural response of DNA minicircles to superhelical stress, whereas the coarse-grained simulations extend conformational sampling and reveal the broader ensemble of topological states accessible on longer timescales, as visualized in the Supplementary movie SM2

### MD simulations elucidate conformational dynamics and writhing of DNA homo- and hetero catenanes

DNA catenanes possess an additional topological constraint arising from the mechanical interlocking of two circular DNA duplexes [32]. These structures occur naturally as intermediates in many biological processes [3, 6, 66], and have also emerged as important building blocks in DNA nanotechnology owing to their unique topological and mechanical properties [13, 14, 67, 68]. To investigate how this topological constraint influences DNA conformational dynamics, we created AA and Martini CG models of homo (100-100 bps) and heterocatenanes (100-200 bps) in the absence of any superhelical stress, Table 1 and performed MD simulations for 0.5*µ*s and 10*µ*s using AMBER and GROMACS, respectively Supplementary movie SM3.

Visualization of instantaneous snapshots from AA simulations shows that both homo- and hetero-catenanes maintain their overall conformational integrity, with only small local structural fluctuations throughout the 0.5*µ*s dynamics (Figure 4 A, B). Homocatenane exhibits cooperative conformational fluctuations of the two 100 bp rings while retaining a compact relative arrangement between them (Figure 4 A). In contrast, heterocatenane displays asymmetric structural dynamics, with the larger 200 bp ring undergoing greater conformational deformation than the smaller 100 bp ring (Figure 4 B). Chain-wise RMSD and *R_g_* analyses further support these observations (Supplementary Figure S8).

**Figure 4.**
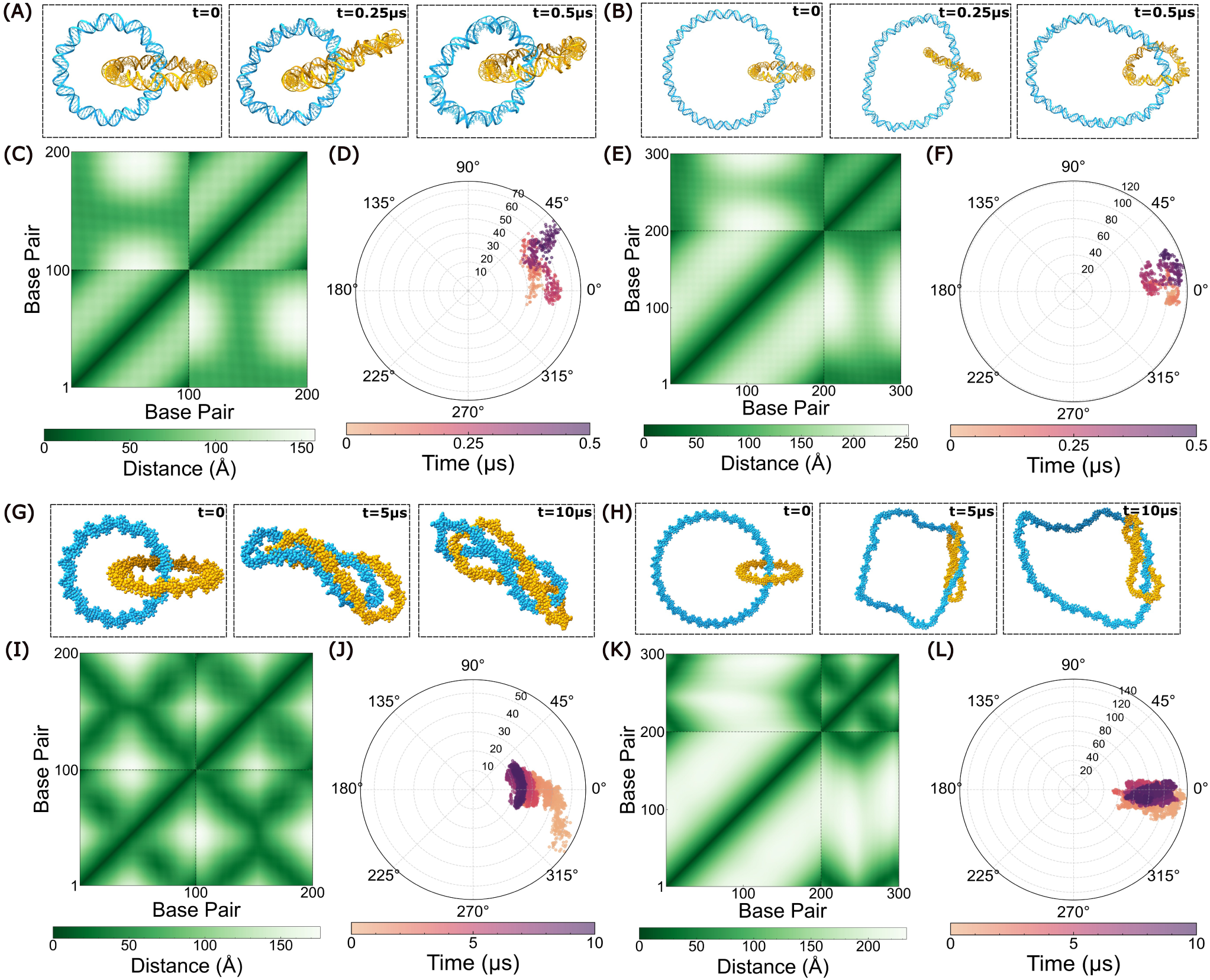
Multiscale structural characterization and inter-ring dynamics of homo and hetero DNA catenanes. Instantaneous snapshots of (A) homocatenane (100 bp + 100 bp) and (B) heterocatenane (200 bp + 100 bp) at 0, 250 and 0.5*µ*s in AA simulations, water and ions are not shown for clarity. (C, E) Time-averaged base-pair distance matrices (⟨*D*⟩) for the homo- and heterocatenanes obtained from the AA simulations. (D, F) Relative rotational dynamics of the interlocked DNA rings represented by the rotational displacement angle (*ϕ*) as a function of simulation time for the homo- and heterocatenanes, respectively. (G, H) Instantaneous snapshots of the homo- and heterocatenanes at 0 5 and 10 *µ*s during the CG simulations. (I, K) Time-averaged base-pair distance matrices obtained from the CG simulations. (J, L) Relative rotational dynamics of the interlocked DNA rings from the CG simulations, showing the evolution of the rotational displacement angle (*ϕ*) over the simulation trajectory.

The time–averaged distance matrix ⟨*D*⟩ between the base-pair COM and the rotational displacement angle (*ϕ*) between the two DNA rings reveal distinct contact patterns and relative orientations (Figure 4 C–F). The definitions of these structural descriptors are provided in Supplementary Section S2 (A and D) and reported earlier by Song *et al.* for DNA catenanes [32]. Homo- and heterocatenane display a symmetric interaction pattern with persistent inter-ring contacts evident from the off-diagonal blocks (Figure 4 C,E), respectively. The rotational displacement (*ϕ*) analysis further shows that homocatenane exhibits a gradual directional rotation of one ring around the other (Figure 4 D). Heterocatenane on the other hand, samples a narrower range of rotational angles, indicating restricted rotational flexibility of one ring around the other.(Figure 4 F).

The modest fluctuations in ΔTw and ΔWr for each DNA ring reveal the weak supercoiling effects seen across both homo and heterocatenanes (Supplementary Figures S9, S10). The free energy landscapes (FELs) obtained using the Boltzmann inversion of the probability distribution of the COM distance between the rings (d*_COM_*) and the relative twist angle (*θ*_twist_) further corroborate the topological dynamics of the system (Supplementary Figure S11 A,B). We observe localized low free-energy basins, centered around the initial conformational states, suggesting a limited exploration of the topological possibilities at the AA simulation timescale. The Na^+^ ion density distributions (Supplementary Figures S12 A, S13 A) reveal preferential localization of counterions around the negatively charged DNA backbone of both rings.

We simulated 10*µ*s conformational dynamics of DNA catenanes using the Martini CG model (Figure 4 G,H). Homocatenane exhibits a pronounced global rearrangement, with both rings adopting elongated and folded conformations (Figure 4 G). Similarly, heterocatenane displays structural deformation of the 200 bp ring, whereas the 100 bp ring remains highly compact and writhed within the catenation region (Figure 4 H). Chain-wise RMSD and *R_g_* analyses further support these large-scale structural rearrangements observed during the CG simulations. (Supplementary Figure S14).

Compared to AA simulations, the time–averaged base-pair distance matrices ⟨*D*⟩ exhibit more extensive off-diagonal contact patterns, consistent with enhanced conformational sampling and supercoiling effects over the extended simulation timescale (Figure 4 I,K). The rotational (*ϕ*) analysis elucidates the relative orientation dynamics of the interlocked rings throughout the trajectories, (Figure 4 J,L). Homocatenane undergoes progressive directional rotation between the two rings during the initial half of the simulation, before both rings simultaneously occupy a writhed conformation which restricts the rotational motion of one ring around another. This causes the relative rotational angle (*ϕ*) to hover around very small values in the later half of the simulation (Figure 4 J). Interestingly in the case of heterocatenane, the system samples a much narrower range of rotational states, due to rapid writhing of the smaller ring, which in turn blocks it’s relative angular rotation around the 200bp ring, causing the angle (*ϕ*) to be bounded to very small values (Figure 4 L).

During longer time-scale CG simulations, we predominantly observe strong negative writhing in both the rings of homocatenane setup (Supplementary Figure S9). Interestingly we find an asymmetric supercoiling behaviour for heterocatenane (Supplementary Figure S10), where the smaller ring (100bp) undergoes strong negative writhing, but the bigger ring (200bp) remains in a mildly positive writhed state. This writhing of smaller ring leads to persistent off-diagonal contacts observed in the top-right corner of Figure 4K. The large amplitude twist and writhe fluctuations enable the rings to access a wide range of topological conformations, which is reflected in the corresponding 2-D free energy profiles (Supplementary Figure S11 C,D). Unlike the localized free energy basins observed in the AA simulations, the global free energy minima correspond to more compact and highly writhed states. The Na^+^ ion density map reveals a diffuse distribution around the DNA phosphate backbone, consistent with the reduced electrostatic resolution and ion representation in the Martini2 coarse-grained force field (Supplementary Figures S12 B, S13 B).

These conformational variations reveal the structural evolution of the catenated topology across multiple time-scales. AA simulations illustrate the local stable structure due to non-bonded interactions between the DNA rings, while CG simulations allow longer time-scale exploration yielding an alternative, highly supercoiled arrangement of the rings, as visualized in the Supplementary movie SM3. We also observe the effect of relative size difference between the rings, leading to an asymmetric supercoiling behaviour of either rings in the same topologically linked setup.

### Cooperative structural dynamics of DNA Borromean rings

Borromean rings are the simplest example of a wide category of well-studied Brunnian links and have been realized in molecular systems using supramolecular chemistry [22]. They are fundamentally different than a catenane, because in this arrangement any chosen pair of rings is not interlocked, but all 3 rings collectively are topologically restrained. It has also been shown that perfect circles mathematically can never assemble into Borromean rings setup [69] and hence to maintain the structure, they are bound to have some degree of eccentricity. Although Seeman and coworkers demonstrated the synthesis of DNA Borromean rings [17] about three decades ago, so far no computational studies have tried to elucidate their nanoscale dynamics. We employed `SDNA`, for the first time, to create AA and CG models of DNA Borromean rings made up of 120bp × 3, as described in the methods section. Multiscale simulations were carried out for 0.5*µ*s and 10*µ*s in AA and CG resolution respectively to reveal their structural evolution at the molecular level.

Instantaneous snapshots from the multiscale simulation show a glimpse of the dynamics of the three DNA rings in Borromean topology (Figure 5A, B), the complete dynamics is illustrated in Supplementary movie SM4. In the initial configuration, the three DNA rings are arranged orthogonal to one another, with COMs of all three rings coinciding at the same point in 3D space, resulting in an average pairwise COM distance of ⟨*d*_COM_⟩ = 0. During the dynamics, the COMs of the individual rings gradually drift apart, giving rise to finite pairwise COM distances while preserving the characteristic mutually orthogonal arrangement of the Borromean topology. Ring-wise RMSD and *R_g_* analyses show that all DNA rings are showing different dynamics and compacting differently from the initial conformation in both AA and CG simulations, (Supplementary Figure S15).

**Figure 5.**
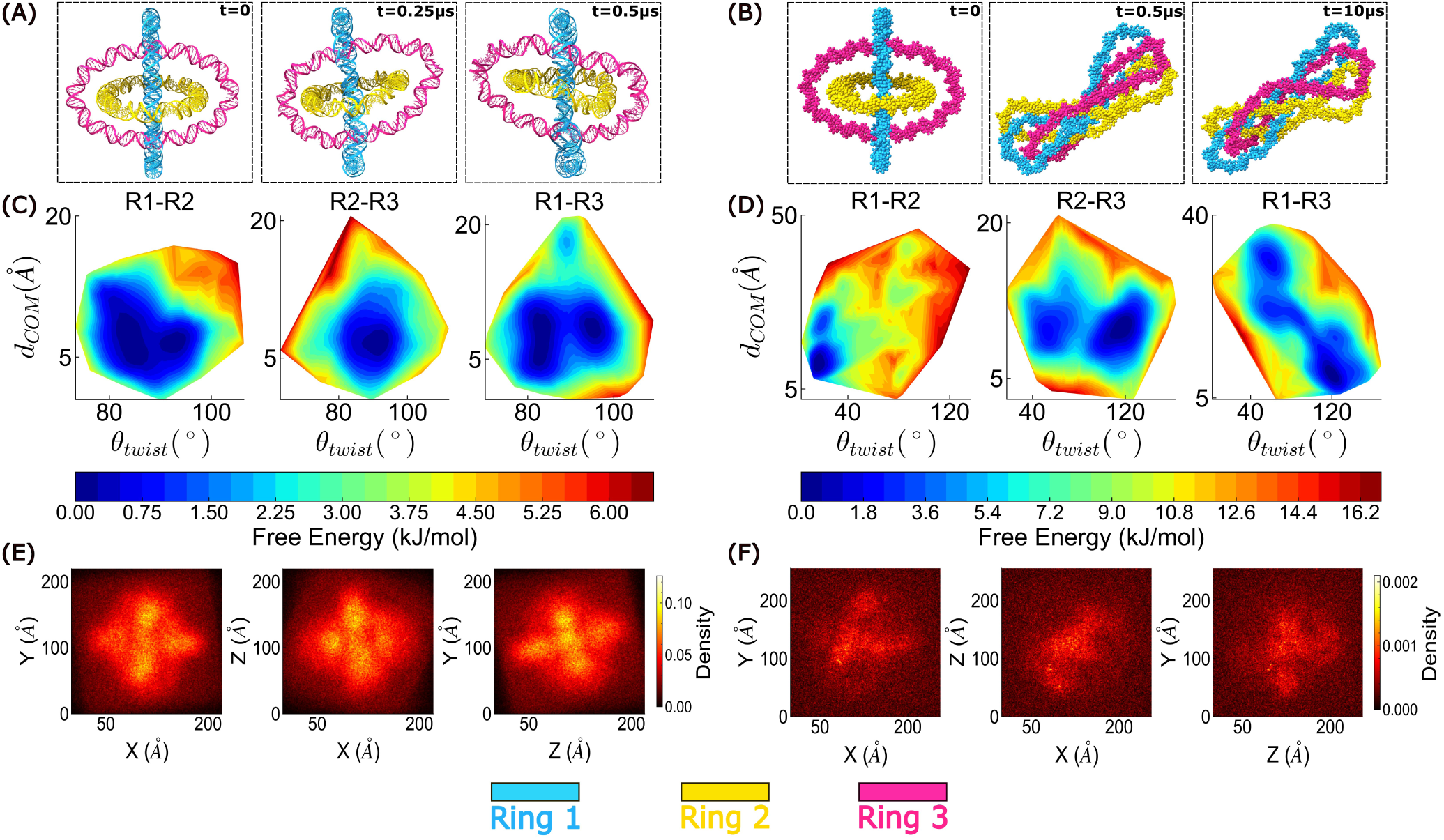
Structural organization and dynamic behavior of DNA Borromean rings from multiscale simulations. (A, B) Representative snapshots of the DNA Borromean rings at different simulation time points from the AA and CG simulations, respectively. (C, D) 2-D free energy landscapes projected onto the pairwise center-of-mass distance (*d_COM_*) and relative twist angle (*θ*_twist_) between the rings, obtained from the AA and CG simulation trajectories, respectively. (E, F) Two-dimensional projections of the Na^+^ ion density surrounding the DNA backbone in three orthogonal planes from the AA and CG simulations, respectively.

To characterize the relative organization of the interlocked rings, the 2-D free energy profile was visualized using the pairwise COM distance 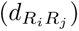 and the relative twist angle 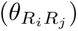 between each pair of rings (*R_i_* and *R_j_*) as the collective variable (Figure 5 C,D). The AA simulation exhibits a single, well-defined free energy minimum for all three pairs, (Figure 5C). This indicates that the Borromean assembly retains a preferred relative orientation while undergoing only limited thermal fluctuations throughout the 0.5*µ*s AA simulation.

In contrast, the CG simulation samples substantially broader free energy basins spanning across larger COM separations and a wider distribution of relative twist angles. The enhanced conformational space explored over the 10*µ*s trajectory reflects the increased flexibility achieved in the CG simulations (Figure 5 D). Notably, the three ring pairs exhibit distinct free energy landscapes rather than identical minima, indicating that each pair experiences a unique local mechanical environment despite the topological equivalence of the Borromean link. These asymmetric free energy profiles highlight the heterogeneous mechanical constraints imposed by the collective three-ring topology and demonstrate the ability of the CG simulations to capture wider phase space while preserving the overall topological integrity of the Borromean structure.

The 2-D free energy plots corroborated by the visualization of simulation trajectories, suggest that mutually aligned plectonemic arrangement are favoured more than the roughly orthogonal arrangement sampled in the initial ∼ 1*µs* of the AA simulations. In these mutually aligned structures, there exist multiple metastable intermediates corresponding to different values of *COM* distances between each ring in the Borromean topology. The time-averaged base-pair distance matrices 〈D〉 co-relates with these findings (Supplementary Figure S16). The AA matrix displays smooth intra- and inter-ring distance gradients consistently, while the distance matrix in CG simulations shows off-diagonal pattern of alternating contact and separation regions, reflecting the formation of plectoneme-like structural rearrangements between the three rings over the longer simulation timescale.

Twist and writhe analyses show that the AA simulations exhibit only minor topological fluctuations while preserving the initial Borromean arrangement, with all three rings displaying weak positive supercoiling. In contrast, the longer CG simulations capture the emergence of negatively supercoiled plectonemes, resulting in a tilted ring arrangement with sustained inter- and intra-ring contacts (Supplementary Figure S17). Further, we observed starkly different twist-writhe partitioning profiles for each individual ring in the CG simulations. Ring 1 exhibits only a weak writhing response, while Ring 3 undergoes a modest accumulation of negative writhe. By contrast, Ring 2 consistently develops significantly greater negative writhe that remains stable over the course of the simulation. This asymmetric yet coordinated redistribution of writhe highlights the cooperative coupling between supercoiling-induced torsional stress and the topological constraints inherent to the Borromean linkage.

The AA simulation of DNA Borromean rings exhibit a pronounced density of Na^+^ ion atmosphere that closely follows the elliptical geometry of the rings, with enhanced ion accumulation near the negatively charged DNA backbone (Figure 5E). In contrast, the CG simulations exhibit a markedly altered ion distribution characterized by sparse, localized density hotspots and the absence of a continuous counterion cloud surrounding the DNA (Figure 5F).

Overall, the structural and mechanical analyses from the multiscale simulations reveal different dynamical behaviors in the AA and CG models, as visualized in the Supplementary movie SM4. While the AA simulations capture only modest conformational and topological fluctuations, the extended CG simulations uncover large-scale plectoneme-like rearrangements, heterogeneous redistribution of writhe, and a substantially broader conformational free-energy landscape. These results demonstrate the complementary strengths of the two simulation approaches in characterizing the dynamics of Borromean DNA assemblies.

## Conclusion

Since the dawn of 21st centaury, computational modeling tools have significantly advanced DNA nanotechnology or nucleic acid research in general by enabling the experimental design [38] and simulations of increasingly complex DNA systems [37, 39, 43, 46, 47]. However,the construction of user-defined 3D DNA topologies often requires specialized expertise, multiple software packages, and extensive manual preprocessing before molecular simulations can be performed. As DNA nanotechnology evolves from proof-of-concept structures in test tubes to functional nanostructures with real world applications, there is a growing need for a computational framework that optimize the design by multiscale structural modeling and simulation of complex DNA nanostructures. While the ergodicity assumption implies that sufficiently long MD simulations should sample the complete conformational ensemble irrespective of the initial structure, in practice, the predicted conformational dynamics often depend on the starting 3D model due to limited simulation timescales.

Here, we present SketchDNA (`SDNA`), an intuitive sketch-to-structure framework that enables users to design 3D DNA geometries and automatically convert them into AA and Martini CG ds-DNA structures. Built upon the MDNA framework [37], `SDNA` is an open-source platform that is freely accessible at https://sdna.biotech.iith.ac.in. The platform enables an intuitive and hassle-free generation of diverse DNA architectures with user-defined nucleotide sequences, topologies, and superhelical densities, while providing interactive visualization of the resulting AA and CG structures directly within the web interface. To demonstrate the simple computational pipeline of `SDNA`, we created three representative topological DNA architectures, including minicircles, catenanes, and Borromean rings, and characterized their structural and topological behavior through multiscale MD simulations.

The asymmetric mechanical response of DNA minicircle to torsional stress was reflected in the distinct twist– writhe partitioning observed for positively and negatively supercoiled minicircles, together with stress-induced DNA kinking in the supercoiled states. These observations are consistent with previous experimental studies and multiscale simulations of DNA minicircles [4, 57, 58, 59, 60, 70, 71]. Multiscale simulations of homo- and heterocatenanes reproduced the characteristic structural dynamics and topological behavior of interlocked DNA rings reported in previous atomistic studies [32]. While previous investigations have been built on atomistic [32] and specific CG models [72], we extend this work by performing integrated and generalized Martini CG simulations of DNA catenanes to explore their structural dynamics at the molecular level. Although DNA Borromean rings were synthesized by Ned Seeman’s group in 1997, no molecular simulation study of this iconic topological architecture has been reported to date; a gap that reflects the absence of accessible, user-friendly modeling tools such as `SDNA`. The multiscale simulations of the DNA Borromean ring characterize, for the first time, the equilibrium structure and dynamics of three interlocked DNA rings in a Borromean topology.

In total, `SDNA` provides a simple yet powerful multiscale computational framework for investigating the structure, stability, and conformational dynamics at atomistic and coarse-grained resolutions in topological DNA systems. The proof-of-concept simulations illustrate the utility of the toolkit. While atomistic models capture detailed local structural fluctuations, coarse-grained simulations efficiently explore long-timescale conformational and topological rearrangements, highlighting the complementary strengths of the two resolutions.

`SDNA` is an open-source toolkit with its source code available on GitHub at https://github.com/sudhishgupta/ SDNA, together with documentation, installation instructions, a step-by-step user guide, and a description of the complete pipeline. It provides a unified and accessible platform that bridges intuitive geometric design with multiscale molecular modeling, making the computational investigation of complex DNA topologies faster, easier, and more accessible to interdisciplinary DNA research. The topological models obtained from `SDNA` can be easily tailored to protein-DNA interactions and elucidate the mechanistic interplay between DNA topology, gene regulation, and cellular function. Future developments will focus on expanding `SDNA` to support a broader range of nucleic acid architectures, including branched DNA nanostructures and hybrid DNA/RNA systems, integrating advanced structural and simulation analysis tools into a unified workflow, *etc. `SDNA`* is expected to enable the rational design, multiscale modeling, and mechanistic investigation of increasingly complex DNA-based synthetic biomolecular assemblies. The simple and open access pipeline of our toolkit will broaden the usage of multiscale modeling of topological DNA systems.

## Funding

PY acknowledges the Council of Scientific and Industrial Research (CSIR), India, for the doctoral fellowship. This work was supported by the Seed Research Grant (SG124) from IIT Hyderabad (HJ), the INSPIRE Faculty Fellowship (IFA-20-PH-256) (HJ), and the Science and Engineering Research Board (SERB), India, through Grant No. SRG/2022/002109 (HJ).

## Acknowledgements

The authors thank Dr. Mahipal Ganji (IISc Bangalore) and Dr. Kush Coshic (Max Planck Institute of Biophysics) for the helpful discussion. The authors acknowledge the support of the Center for Development of Advanced Computing (CDAC), Pune, and IIT Hyderabad for providing access to the PARAM Seva supercomputing facility under the National Supercomputing Mission (NSM).

## Supplementary Data

Supplementary data, including detailed methods, simulation analyses, figures, and discussion, are available online in a consolidated Supporting Information file. A video tutorial explaining the step-by-step usage of the `SDNA` tool is provided as supplementary movie SM1. Three simulation movies illustrating the all-atom and coarse-grained MD simulation trajectories of the systems are also available online. The captions for these movies are provided in the Supporting Information.

## Data Availability

The topology and initial coordinate files for AMBER all-atom simulations and coarse-grained Martini models in GROMACS format, input simulation files, along with the pipeline for conducting all the analyses, are available on GitHub at https://github.com/sudhishgupta/SDNA

## Conflict of interest statement

None declared.

## Notes

https://sdna.biotech.iith.ac.in/

https://github.com/sudhishgupta/SDNA

### Competing Interest Statement

The authors have declared no competing interest.

### Summary of Updates

This version of the manuscript has been revised to correct minor grammatical and formatting errors.

## References

1. James D Watson and Francis HC Crick. Molecular structure of nucleic acids: a structure for deoxyribose nucleic acid. Nature, 171(4356):737–738, 1953.

2. Shuai Li, Charan Vemuri, and Chongyi Chen. DNA topology: A central dynamic coordinator in chromatin regulation. Current Opinion in Structural Biology, 87:102868, 2024.

3. Yves Pommier, Yilun Sun, Shar-yin N Huang, and John L Nitiss. Roles of eukaryotic topoisomerases in transcription, replication and genomic stability. Nature reviews Molecular cell biology, 17(11):703–721, 2016.

4. Rossitza N Irobalieva, Jonathan M Fogg, Daniel J Catanese Jr, Thana Sutthibutpong, Muyuan Chen, Anna K Barker, Steven J Ludtke, Sarah A Harris, Michael F Schmid, Wah Chiu, et al. Structural diversity of supercoiled DNA. Nature Communications, 6(1):8440, 2015.

5. Radost Waszkiewicz, Maduni Ranasinghe, Jonathan M Fogg, Daniel J Catanese Jr, Maria L Ekiel-Jeżewska, Maciej Lisicki, Borries Demeler, Lynn Zechiedrich, and Piotr Szymczak. DNA supercoiling-induced shapes alter minicircle hydrodynamic properties. Nucleic Acids Research, 51(8):4027–4042, 2023.

6. David E Adams, Eugene M Shekhtman, E Lynn Zechiedrich, Molly B Schmid, and Nicholas R Cozzarelli. The role of topoisomerase iv in partitioning bacterial replicons and the structure of catenated intermediates in DNA replication. Cell, 71(2):277–288, 1992.

7. Sneha Shahu, Sreeja Baira, Sudhish Gupta, and Mahipal Ganji. A DNA-binding protein senses DNA superhelicity to switch between bridging and nucleoprotein filament formation. bioRxiv, 2026. doi: 10.64898/2026.05.23.727375.

8. Xiaomeng Jia, Xiang Gao, Shuming Zhang, James T. Inman, Yifeng Hong, Anupam Singh, Fahad Rashid, James M. Berger, Smita S. Patel, and Michelle D. Wang. Torsion is a dynamic regulator of DNA replication stalling and reactivation. Nature Communications, 16, 2025. doi: 10.1038/s41467-025-65567-5.

9. Nadrian C Seeman and Hanadi F Sleiman. DNA nanotechnology. Nature Reviews Materials, 3(1):17068, 2017.

10. Paul WK Rothemund. Folding DNA to create nanoscale shapes and patterns. Nature, 440(7082):297–302, 2006.

11. Dhiraj Bhatia, Senthil Arumugam, Michel Nasilowski, Himanshu Joshi, Christian Wunder, Valérie Chambon, Ved Prakash, Chloé Grazon, Brice Nadal, Prabal K Maiti, et al. Quantum dot-loaded monofunctionalized DNA icosahedra for single-particle tracking of endocytic pathways. Nature Nanotechnology, 11(12):1112–1119, 2016.

12. Thomas Thibault, Jeril Degrouard, Patrick Baril, Chantal Pichon, Patrick Midoux, and Jean-Marc Malinge. Production of DNA minicircles less than 250 base pairs through a novel concentrated DNA circularization assay enabling minicircle design with nf-*κ*b inhibition activity. Nucleic Acids Research, 45(5):e26–e26, 2017.

13. Thorsten L Schmidt and Alexander Heckel. Construction of a structurally defined double-stranded DNA catenane. Nano letters, 11(4):1739–1742, 2011.

14. Zai-Sheng Wu, Zhifa Shen, Kha Tram, and Yingfu Li. Engineering interlocking DNA rings with weak physical interactions. Nature Communications, 5(1):4279, 2014.

15. Dongran Han, Suchetan Pal, Yan Liu, and Hao Yan. Folding and cutting DNA into reconfigurable topological nanostructures. Nature Nanotechnology, 5(10):712–717, 2010.

16. Damian Ackermann, Thorsten L Schmidt, Jeffrey S Hannam, Chandra S Purohit, Alexander Heckel, and Michael Famulok. A double-stranded DNA rotaxane. Nature Nanotechnology, 5(6):436–442, 2010.

17. Chengde Mao, Weiqiong Sun, and Nadrian C Seeman. Assembly of borromean rings from DNA. Nature, 386 (6621):137, 1997.

18. David Winogradoff, Pin-Yi Li, Himanshu Joshi, Lauren Quednau, Christopher Maffeo, and Aleksei Aksimentiev. Chiral systems made from DNA. Advanced Science, 8(5):2003113, 2021.

19. R. Oliynyk and George M. Church. Circular vectors as an efficient, fully synthetic, cell-free approach for preparing small circular DNA as a plasmid substitute for guide RNA expression in crispr–cas9 genome editing. Nature Protocols, 20:2942 –2959, 2025. doi: 10.1038/s41596-024-01138-0.

20. Meng Liu, Qiang Zhang, Zhongping Li, Jimmy Gu, John D Brennan, and Yingfu Li. Programming a topologically constrained DNA nanostructure into a sensor. Nature Communications, 7(1):12074, 2016.

21. Hendrik Dietz, Shawn M Douglas, and William M Shih. Folding DNA into twisted and curved nanoscale shapes. Science, 325(5941):725–730, 2009.

22. Kelly S. Chichak, Stuart J. Cantrill, Anthony R. Pease, Sheng-Hsien Chiu, Gareth W. V. Cave, Jerry L. Atwood, and J. Fraser Stoddart. Molecular borromean rings. Science, 304:1308–1312, 2004. doi: 10.1126/science.1096914.

23. Mahipal Ganji, Indra A Shaltiel, Shveta Bisht, Eugene Kim, Ana Kalichava, Christian H Haering, and Cees Dekker. Real-time imaging of DNA loop extrusion by condensin. Science, 360(6384):102–105, 2018.

24. Abhinav Banerjee, Saanya Yadav, Vedanth Shree Vidwath, Simanta Kalita, Sarit S Agasti, Himanshu Joshi, and Mahipal Ganji. Proximity-based super-resolution imaging enabled by DNA base-stacking interactions. Small, page e07139, 2025.

25. Stephanie Johnson, Martin Lindén, and Rob Phillips. Sequence dependence of transcription factor-mediated DNA looping. Nucleic Acids Research, 40(16):7728–7738, 2012.

26. Kush Coshic, Christopher Maffeo, David Winogradoff, and Aleksei Aksimentiev. The structure and physical properties of a packaged bacteriophage particle. Nature, 627(8005):905–914, 2024.

27. Himanshu Joshi and Prabal K Maiti. Structure and electrical properties of DNA nanotubes embedded in lipid bilayer membranes. Nucleic Acids Research, 46(5):2234–2242, 2018.

28. Himanshu Joshi, Atul Kaushik, Nadrian C Seeman, and Prabal K Maiti. Nanoscale structure and elasticity of pillared DNA nanotubes. ACS Nano, 10(8):7780–7791, 2016.

29. Jejoong Yoo, Sangwoo Park, Christopher Maffeo, Taekjip Ha, and Aleksei Aksimentiev. DNA sequence and methylation prescribe the inside-out conformational dynamics and bending energetics of DNA minicircles. Nucleic Acids Research, 49(20):11459–11475, 2021.

30. Matthew Burman and Agnes Noy. Atomic description of the reciprocal action between supercoils and melting bubbles on linear DNA. Physical Review Letters, 134(3):038403, 2025.

31. Himanshu Joshi, Anjan Dwaraknath, and Prabal K Maiti. Structure, stability and elasticity of DNA nanotubes. Physical Chemistry Chemical Physics, 17(2):1424–1434, 2014.

32. Yeonho Song, Minjung Kim, Bong June Sung, and Jun Soo Kim. Computational characterization of DNA catenanes. Journal of Chemical Theory and Computation, 21(19):9967–9981, 2025.

33. Siewert J Marrink, H Jelger Risselada, Serge Yefimov, D Peter Tieleman, and Alex H De Vries. The martini force field: coarse-grained model for biomolecular simulations. The Journal of Physical Chemistry B, 111(27): 7812–7824, 2007.

34. Thomas E Ouldridge, Petr Šulc, Flavio Romano, Jonathan PK Doye, and Ard A Louis. DNA hybridization kinetics: zippering, internal displacement and sequence dependence. Nucleic Acids Research, 41(19):8886–8895, 2013.

35. Thomas J Macke and David A Case. Modeling unusual nucleic acid structures. ACS Publications, 1998.

36. Shuxiang Li, Wilma K Olson, and Xiang-Jun Lu. Web 3DNA 2.0 for the analysis, visualization, and modeling of 3d nucleic acid structures. Nucleic Acids Research, 47(W1):W26–W34, 2019.

37. Thor van Heesch, Enrico Skoruppa, Peter G Bolhuis, Helmut Schiessel, and Jocelyne Vreede. MDNA: a software module for DNA structure generation and analysis. Nucleic Acids Research, 54(10):gkag549, 2026.

38. Shawn M Douglas, Adam H Marblestone, Surat Teerapittayanon, Alejandro Vazquez, George M Church, and William M Shih. Rapid prototyping of 3d DNA-origami shapes with caDNAno. Nucleic Acids Research, 37 (15):5001–5006, 2009.

39. Sean Williams, Kyle Lund, Chenxiang Lin, Peter Wonka, Stuart Lindsay, and Hao Yan. Tiamat: a three-dimensional editing tool for complex DNA structures. In International workshop on DNA-based computers, pages 90–101. Springer, 2008.

40. Do-Nyun Kim, Fabian Kilchherr, Hendrik Dietz, and Mark Bathe. Quantitative prediction of 3d solution shape and flexibility of nucleic acid nanostructures. Nucleic Acids Research, 40(7):2862–2868, 2012.

41. Erik Benson, Abdulmelik Mohammed, Johan Gardell, Sergej Masich, Eugen Czeizler, Pekka Orponen, and Björn Högberg. DNA rendering of polyhedral meshes at the nanoscale. Nature, 523(7561):441–444, 2015.

42. Rémi Veneziano, Sakul Ratanalert, Kaiming Zhang, Fei Zhang, Hao Yan, Wah Chiu, and Mark Bathe. Designer nanoscale DNA assemblies programmed from the top down. Science, 352(6293):1534–1534, 2016.

43. Elisa De Llano, Haichao Miao, Yasaman Ahmadi, Amanda J Wilson, Morgan Beeby, Ivan Viola, and Ivan Barisic. Adenita: interactive 3d modelling and visualization of DNA nanostructures. Nucleic Acids Research, 48(15):8269–8275, 2020.

44. Wolfgang G Pfeifer, Chao-Min Huang, Michael G Poirier, Gaurav Arya, and Carlos E Castro. Versatile computer-aided design of free-form DNA nanostructures and assemblies. Science Advances, 9(30):eadi0697, 2023.

45. Antti Elonen, Leon Wimbes, Abdulmelik Mohammed, and Pekka Orponen. DNAforge: a design tool for nucleic acid wireframe nanostructures. Nucleic Acids Research, 52(W1):W13–W18, 2024.

46. Christopher Maffeo and Aleksei Aksimentiev. MrDNA: a multi-resolution model for predicting the structure and dynamics of DNA systems. Nucleic Acids Research, 48(9):5135–5146, 2020.

47. Erik Poppleton, Roger Romero, Aatmik Mallya, Lorenzo Rovigatti, and Petr Šulc. OxDNA. org: a public webserver for coarse-grained simulations of DNA and RNA nanostructures. Nucleic Acids Research, 49(W1): W491–W498, 2021.

48. Himanshu Joshi, Dhiraj Bhatia, Yamuna Krishnan, and Prabal K Maiti. Probing the structure and in silico stability of cargo loaded DNA icosahedra using md simulations. Nanoscale, 9(13):4467–4477, 2017.

49. Himanshu Joshi, Chen-Yu Li, and Aleksei Aksimentiev. All-atom molecular dynamics simulations of membrane-spanning DNA origami nanopores. In DNA and RNA Origami: Methods and Protocols, pages 113–128. Springer, 2023.

50. Djurre H De Jong, Gurpreet Singh, WF Drew Bennett, Clement Arnarez, Tsjerk A Wassenaar, Lars V Schafer, Xavier Periole, D Peter Tieleman, and Siewert J Marrink. Improved parameters for the martini coarse-grained protein force field. Journal of chemical theory and computation, 9(1):687–697, 2013.

51. Jaakko J. Uusitalo, Helgi I. Ingólfsson, Parisa Akhshi, D. Peter Tieleman, and Siewert J. Marrink. Martini coarse-grained force field: Extension to DNA. Journal of Chemical Theory and Computation, 11(8):3932–3945, 2015. doi: 10.1021/acs.jctc.5b00286.

52. David A Case, Hasan Metin Aktulga, Kellon Belfon, David S Cerutti, G Andrés Cisneros, Vinícius Wilian D Cruzeiro, Negin Forouzesh, Timothy J Giese, Andreas W Gotz, Holger Gohlke, et al. Ambertools. Journal of chemical information and modeling, 63(20):6183–6191, 2023.

53. Mark James Abraham, Teemu Murtola, Roland Schulz, Szilárd Páll, Jeremy C Smith, Berk Hess, and Erik Lindahl. Gromacs: High performance molecular simulations through multi-level parallelism from laptops to supercomputers. SoftwareX, 1:19–25, 2015.

54. James C Phillips, David J Hardy, Julio DC Maia, John E Stone, João V Ribeiro, Rafael C Bernardi, Ronak Buch, Giacomo Fiorin, Jérôme Hénin, Wei Jiang, et al. Scalable molecular dynamics on cpu and gpu architectures with namd. The Journal of Chemical Physics, 153(4), 2020.

55. Marie Zgarbová, Jiri Sponer, and Petr Jurecka. Z-DNA as a touchstone for additive empirical force fields and a refinement of the alpha/gamma DNA torsions for amber. Journal of Chemical Theory and Computation, 17 (10):6292–6301, 2021.

56. Sergei M Mirkin. DNA topology: Fundamentals. Encyclopedia of Life Sciences, 2001. doi: 10.1038/npg.els.0001038.

57. Filip Lankaš, Richard Lavery, and John H Maddocks. Kinking occurs during molecular dynamics simulations of small DNA minicircles. Structure, 14(10):1527–1534, 2006.

58. Troy A Lionberger, Davide Demurtas, Guillaume Witz, Julien Dorier, Todd Lillian, Edgar Meyhöfer, and Andrzej Stasiak. Cooperative kinking at distant sites in mechanically stressed DNA. Nucleic Acids Research, 39(22):9820–9832, 2011.

59. JS Mitchell, CA Laughton, and Sarah A Harris. Atomistic simulations reveal bubbles, kinks and wrinkles in supercoiled DNA. Nucleic Acids Research, 39(9):3928–3938, 2011.

60. Alice LB Pyne, Agnes Noy, Kavit HS Main, Victor Velasco-Berrelleza, Michael M Piperakis, Lesley A Mitchenall, Fiorella M Cugliandolo, Joseph G Beton, Clare EM Stevenson, Bart W Hoogenboom, et al. Base-pair resolution analysis of the effect of supercoiling on DNA flexibility and major groove recognition by triplex-forming oligonucleotides. Nature Communications, 12(1):1053, 2021.

61. Thana Sutthibutpong, Sarah A. Harris, and Agnes Noy. Comparison of molecular contours for measuring writhe in atomistic supercoiled DNA. Journal of Chemical Theory and Computation, 11(6):2768–2775, 2015. doi: 10.1021/acs.jctc.5b00035.

62. Sarah A Harris, Charles A Laughton, and Tanniemola B Liverpool. Mapping the phase diagram of the writhe of DNA nanocircles using atomistic molecular dynamics simulations. Nucleic Acids Research, 36(1):21–29, 2008.

63. Anupam Mondal and Arnab Bhattacherjee. Torsional behaviour of supercoiled DNA regulates recognition of architectural protein fis on minicircle DNA. Nucleic Acids Research, 50(12):6671–6686, 2022.

64. Mehmet Sayar, Barış Avşaroǧlu, and Alkan Kabakçıoǧlu. Twist-writhe partitioning in a coarse-grained DNA minicircle model. Physical Review E, 81, 2010. doi: 10.1103/PhysRevE.81.041916.

65. Supriyo Naskar and Prabal K Maiti. Mechanical properties of DNA and DNA nanostructures: Comparison of atomistic, martini and oxDNA models. Journal of Materials Chemistry B, 9(25):5102–5113, 2021.

66. Isabelle Lucas, Thomas Germe, Marianne Chevrier-Miller, and Olivier Hyrien. Topoisomerase ii can unlink replicating DNA by precatenane removal. The EMBO journal, 20(22):6509–6519, 2001.

67. Chun-Hua Lu, Alessandro Cecconello, and Itamar Willner. Recent advances in the synthesis and functions of reconfigurable interlocked DNA nanostructures. Journal of the American Chemical Society, 138(16):5172–5185, 2016.

68. Johann Elbaz, Alessandro Cecconello, Zhiyuan Fan, Alexander O Govorov, and Itamar Willner. Powering the programmed nanostructure and function of gold nanoparticles with catenated DNA machines. Nature Communications, 4(1):2000, 2013.

69. Bernt Lindstrom and Hans-Olov Zetterstrom. Borromean circles are impossible. The American Mathematical Monthly, 98:340, 1991. doi: 10.2307/2323803.

70. Quan Du, Alexander Kotlyar, and Alexander Vologodskii. Kinking the double helix by bending deformation. Nucleic Acids Research, 36(4):1120–1128, 2008.

71. Mehmet Sayar, Barış Avşaroǧlu, and Alkan Kabakçıoǧlu. Twist-writhe partitioning in a coarse-grained DNA minicircle model. Physical Review E—Statistical, Nonlinear, and Soft Matter Physics, 81(4):041916, 2010.

72. Zhongyan Zhang, Wenbo Zhao, Zhiyuan Cheng, Guojie Zhang, and Hong Liu. Olympic gels formed through catenation of dsDNA rings regulated by topoisomerase ii: A coarse-grained model. The Journal of Chemical Physics, 160(5), 2024.

