## Supplementary Information for "SketchDNA: A GUI-Enabled Toolkit for Multiscale Modeling of Topological DNA Structures"

### Table of Contents:


**S1. Materials and Methods** …………………………………………………………..………..(3-6)

(A) All-atom model building and simulation protocol …………………………..………..(3-4)

(B) Martini model ………………………………………………………………………….... (4-5)

(C) Simulation protocol for coarse-grained using Martini models in GROMACS ….………(5-6)

**S2. Analysis schemes used for AA and CG simulation trajectories** …………….….. (6-9)

**(A)** Contact Distance Matrix………………………………………………………………….(6)

(B) Ion density …………………………… …………………………… …………………..…(7)

(C) Change in twist and writhe …………………………… …………………………………(7-8)

(D) Center of Mass Distance, Relative Twist and Rotation ……………………………………(8)

(E) Free Energy Landscape …………………………… …………………………………… (8-9)

**S3**. Evolution of RMSD and $R_g$ during multiscale simulation of the DNA minicircle …..…(9-10)

**S4.** H- bonds in AA simulation of DNA minicircle with varying superhelical densities.…(10-11)

**S5.** Structural organization of DNA minicircles revealed by contact distance matrix ......(11-12)

**S6.** $Na^+$ ion density distribution in AA simulation around the DNA minicircle …………..(12-13)

**S7.** $Na^+$ ion distribution in CG simulations around the DNA minicircle …………………......(14)

**S8.** Time evolution of the RMSD and $R_g$ during AA simulation of DNA catenanes ………..(15)

**S9.** Topological fluctuations in DNA homocatenane during multiscale simulations …….(16-17)

**S10.** Topological fluctuations DNA heterocatenane during multiscale simulations …….(18-19)

**S11.** 2D Free Energy of DNA catenanes deduced from multiscale simulations……….......(20)

**S12.** $Na^+$ ion distribution around DNA homocatenane from AA and CG MD simulations…(21)

**S13.** $Na^+$ ion distribution around DNA heterocatenane from AA and CG MD simulation.(21-22)

**S14.** Evolution of the RMSD and $R_g$ during CG simulation of DNA catenane ………..…(22-23)

**S15.** Chain-wise RMSD and radius of gyration analyses of the Borromean rings ……...(23-24)

**S16.** Structural organization of Borromean rings revealed by contact distance matrix ...(24-25)

**S17.** Evolution of writhe and twist in individual rings of the Borromean DNA assembly..(25-26)

**S18. AA and CG Supplementary Movies (SM) (36-38)**

**(A)** Supplementary Movie 1 (SM1) Tutorial video ……..………………………..……………. (27)

(A) Supplementary Movie 2 (SM2) of minicircle…………………………………………… …. (28)

(B) Supplementary Movie 3 (SM3) of homo- and hetero catenane …………………………. (29)

(C) Supplementary Movie 4 (SM4) of the Borromean ring ……………………………… …. (30)

**S19. References** ……………………………..……………………………………..…………..(29-33)

### S1. (A) All-atom model building and simulation protocol

All-atom molecular dynamics (MD) simulations of the topologically distinct DNA systems summarized in Table 1 of the main article were performed using the AMBER24 simulation package (1). The initial configurations of the various DNA topologies were obtained using SketchDNA **(SDNA)**, a GUI-based DNA design tool capable of generating both all-atom and coarse-grained models. **SDNA** is capable of creating all kinds of topological DNA nanostructures. To showcase the utility of our tool, three topological DNA nanostructures were considered in this study: minicircles, catenanes, and Borromean rings. The all-atom models of the respective topological structures were obtained from the server. To make the closed circular DNA, the phosphodiester bond was manually added to connect both ends in xLEaP. The DNA structure was solvated in a cubic TIP3P water box, ensuring a minimum buffer of 14 Å around each system using the xLEaP module (2) of AMBER24. The negative charges of the phosphate backbones were neutralized by adding the appropriate number of $Na^+$ ions by creating a Coulombic grid of 1 Å in xLEaP. To maintain the physiological salt concentration of 150 mM, we added appropriate numbers of $Na^+$ and $Cl^-$ ions.

All solvated systems with 150 mM NaCl concentrations were first subjected to energy minimization for 2000 steps using the conjugate gradient algorithm to remove bad contacts and allow water molecules and ions to reorganize around the DNA. During minimization, harmonic positional restraints of 10 kcal/(mol•Å$^2$). were applied to all DNA atoms. The systems were subsequently heated from 100 to 300 K over 1 ns under constant-volume (NVT) conditions, while reducing the positional restraints on DNA non-hydrogen atoms to 1 kcal/(mol•Å$^2$). Following the heating stage, each system was equilibrated for 5 ns in the isothermal-isobaric (NPT) ensemble at 300 K and 1 atm to achieve the correct solvent density and stabilize the system pressure. During equilibration, harmonic restraints of 1 kcal/(mol•Å$^2$) were maintained on the DNA non-hydrogen atoms. Finally, unrestrained production MD simulations were performed at NPT ensemble (T = 300 K, P = 1 atm), with each trajectory extending to 0.5 μs, to characterize the equilibrium structural and dynamical properties of the topologically distinct DNA systems.

Periodic boundary conditions were applied throughout the simulations. Long-range electrostatic interactions were treated using the particle mesh Ewald (PME) method (3), while short-range electrostatic and van der Waals interactions were evaluated using an 8 Å cutoff. The equations of motion were integrated using a 2 fs time step. DNA was modelled using the

OL21 force field (4), and water molecules were represented by the rigid TIP3P model (5). The non-bonded interactions of monovalent $K^+$ and $Cl^-$ ions were described using the Joung-Cheatham ion parameters (6), and DNA-ion interactions were refined using the CUFIX parameter corrections (7). Temperature was maintained using a Langevin thermostat (8) with a collision frequency of 1 $ps^{-1}$, while pressure was regulated using the Monte Carlo barostat. All hydrogen bonds in water molecules were constrained using the SETTLE method (9) of the SHAKE algorithm, while all other covalent bonds involving hydrogen atoms were constrained using the RATTLE algorithm (10), which allowed us to use an integration time step of 2 fs. All simulations were performed using the CUDA-enabled pmemd.cuda engine (11) on NVIDIA RTX 4080 and Tesla V100 GPU platforms.

Simulation trajectories were visualized in 3D using VMD (12) ChimeraX (13) and analyzed with CPPTRAJ (14). The analysis of simulation data, including root mean square deviation (RMSD), radius of gyration (Rg), was performed using CPPTRAJ, a module of AmberTools. The average 3D ion density distributions of $Na^+$ around the DNA topologies were calculated using the VolMap density module of VMD using a 1 Å grid. The hydrogen bond analysis is performed using a custom Tcl script in VMD. Data visualization was performed using Matplotlib and Seaborn.

**S1. (B) Martini model**

The atomistic structures of all DNA topologies were converted to their coarse-grained (CG) representations automatically by the server, using the updated martinize2 script and the Martini2 forcefield. In this model, DNA is represented using a bead-based framework with an approximate mapping ratio of four heavy atoms per bead. The backbone is described by three beads, corresponding to the phosphate group (BB1) and the sugar moiety (BB2 and BB3), with BB3 representing the 3' terminus of the sugar. Nucleobases are modelled as cyclic side-chain structures attached to the backbone through the SC1 bead, which is connected to BB3. Since all systems considered in this study comprise AT-repeat sequences, adenine residues were represented by four side-chain beads (SC1-SC4), whereas thymine residues were represented by three side-chain beads (SC1-SC3). In double-stranded DNA, the SC2 and SC3 beads participate in complementary base-pairing interactions with the opposing strand. In the CG representation of DNA in the Martini model, standard backbone and sugar beads are represented by Lennard-Jones (LJ) particles with diameters of σ = 0.47 nm and σ = 0.43 nm, respectively. Because nucleobases in double-stranded (ds) DNA are stacked at an interbase distance of only ~0.34 nm, these bead sizes are too large to accurately capture base stacking and the planar geometry of the aromatic bases. Therefore, a smaller bead type, referred to as the tiny (T) bead, with a reduced LJ diameter of $\sigma = 0.32$ nm, is used to represent the nucleobases. Hydrogen-bonding interactions between complementary nucleobases were explicitly tuned to capture the specificity of Watson–Crick pairing. To maintain the canonical dsDNA structure, a soft elastic network was implemented using a force constant of 13 kJ/(mol•nm$^2$) and a cutoff distance of 1.2 nm. For further details regarding the Martini model, its development, and parametrization, the reader is referred to the original references (15). It represents each nucleotide as a set of rigid bodies corresponding to the sugar, phosphate, and base groups. These rigid bodies interact through effective sites that capture backbone connectivity, base stacking, and hydrogen-bonding interactions.

#### S1. (C) Simulation protocol for coarse-grain simulations using Martini-2 DNA forcefields in GROMACS

CG MD simulations were performed using the Martini2 DNA forcefield (15,16) within the GROMACS 2023.3 simulation package (17,18). All systems were solvated in a cubic TIP3P water box with a minimum buffer distance of 20 Å from the solute in all directions to avoid periodic boundary interactions. To prevent freezing artifacts, 10% of the total water molecules were replaced with antifreeze water (WF) molecules. The systems were neutralized by adding an appropriate number of $Na^+$ ions, and a physiological ionic strength of 150 mM was maintained by introducing additional $Na^+$ and $Cl^-$ ions.

Prior to production simulations, all systems were subjected to energy minimization using the steepest-descent algorithm while applying harmonic positional restraints to the DNA with a force constant of 10 kJ/(mol•Å$^2$), which were gradually released during equilibration. The minimized systems were subsequently equilibrated in the NPT ensemble for 50 ns with the DNA restraints maintained, followed by an additional 50 ns of equilibration simulation without restraints. Temperature was maintained at 300 K using the velocity-rescaling thermostat (19) with a coupling constant of 2.0 ps, while pressure was controlled isotropically at 1 bar using the Berendsen barostat (20) during equilibration. After this final production run, simulations were subsequently performed in the NPT ensemble for 10 µs. In this step Parrinello-Rahman barostat (21) was used for maintaining the pressure, with a coupling constant of 3.0 ps and a compressibility of $3 \times 10^{-4}$ bar$^{-1}$. Both equilibration and production simulations were carried out using a 20 fs integration time step.

All Martini simulations were performed using the soft elastic network model. The electrostatic interactions were treated using the reaction-field method with a cutoff distance of 1.1nm and a dielectric constant of 15 (22). The same cutoff was used for the van der Waals interactions with the potential-shift cutoff scheme. The neighbour lists were updated every 10 integration steps using the Verlet cutoff scheme. Periodic boundary conditions were applied in all three dimensions.

### S2. Analysis schemes used for AA and CG simulation trajectories

#### S2. (A) Contact Distance Matrix

To characterize structural rearrangements of the system, we computed a base-pair–resolved distance matrix capturing both intra- and inter-component spatial organization. For each configuration, pairwise distances between all base-pair centers of geometry (COMs) were calculated.

Let **A** denote the total number of base pairs in the system. We define an **A × A** time-dependent distance matrix, **D(t)**, whose elements are given by

$$D_{ij}(t) = |\mathbf{r}_i(t) - \mathbf{r}_j(t)| \qquad \text{------------------} \quad \textbf{(1)}$$

where $\mathbf{r_i(t)}$ is the centre-of-mass position of the $i$-$^{th}$ base pair at time $t$.

Base pairs are indexed sequentially across all components of the system, such that each index is 1∈ 1,…,A  corresponding uniquely corresponds to a specific base pair in a given component.

To characterize persistent contacts throughout the simulation, we compute the time-averaged distance matrix, ⟨D⟩, as

$$\langle D \rangle = \frac{1}{T}\sum_{t=0}^{T} D(t) \qquad \text{------------------} \quad \textbf{(2)}$$

Where **T** is the total simulation time.

### S2. (B) Ion density

$Na^+$ ion distributions around the DNA were quantified by constructing a time-averaged spatial occupancy map. Prior to analysis, all frames were aligned to a common reference frame defined by the DNA structure to remove global translation and rotation. Ion positions were then accumulated on a uniform three-dimensional grid with spacing Δ =1 Å, yielding a spatial map of ion presence over the trajectory.

For each voxel, the reported quantity corresponds to the average number of times ions are found within that region during the simulation:

$$\rho(\mathbf{r}) \propto \frac{1}{N_{\mathrm{frames}}} \sum_{t} \sum_{i=1}^{N_{\mathrm{ion}}} I_{\mathrm{bin}}(\mathbf{r}_i(t)) \quad \text{------------------} \quad \textbf{(3)}$$

Where $r_i(t)$ denotes the position of ion (i) at time (t), and $1_{bin}$ is equal to one if the ion lies within a given grid voxel and zero otherwise. This quantity therefore represents a time-averaged ion occupancy field, indicating how frequently ions visit each region of space.

For visualization, the three-dimensional occupancy map was projected onto each of the xy, yz, and zx planes:

$$\rho_{2D}(x,y) = \sum_{z} \rho(x,y,z)$$
$$\rho_{2D}(y,z) = \sum_{x} \rho(x,y,z)$$
$$\rho_{2D}(z,x) = \sum_{y} \rho(x,y,z) \quad \text{------------------} \quad \textbf{(4)}$$

producing a two-dimensional map that reflects the preferred regions of ion accumulation around the DNA over the full simulation.

### S2. (C) Change in twist and writhe

For all-atom simulations, the time evolution of writhe for a closed circular DNA was calculated using WrLINE (23). For coarse-grained systems, we used an in-house modified version of

WrLINE, in which we replaced the C1' atom with the BB2 bead for all calculations. To better illustrate the changes in twist-writhe, we calculated ΔTw and ΔWr, defined as follows:

$$\Delta Tw(t) = Tw|_t - Tw|_{t=0} \qquad \text{------------------} \quad \textbf{(5)}$$

$$\Delta Wr(t) = Wr|_t - Wr|_{t=0} \qquad \text{------------------} \quad \textbf{(6)}$$

**S2. (D) Center of Mass Distance, Relative Twist and Rotation**

The length of the mechanical bond formed by a pair of topologically interlocked rings was quantified by measuring the Center of Mass (COM) distance between two rings $R_i$ and $R_j$ at any given time point $'t'$ as :

$$d_{COM} = COM(R_i)(t) - COM(R_j)(t) \quad \text{------------------} \quad \textbf{(7)}$$

To quantify relative twist and rotation between a pair of topologically interlocked rings, for example, catenane/Borromean links, we defined a normal vector $\mathbf{\tilde{n}_R}$ for each ring $R$, which is defined along the axis perpendicular to the best-fitting plane of the ring, determined from the eigenvector with the smallest principal moment of inertia (24) . The unit normal vector is then obtained as

$$\mathbf{\hat{n_r}} = \frac{\mathbf{\tilde{n_r}}}{|\mathbf{\tilde{n_r}}|} \quad \text{------------------} \quad \textbf{(8)}$$

The relative twist between two rings $R_i$ and $R_j$ is then defined as the angle between the unit normal vector of the rings and can be calculated as

$$\theta_{i,j}^{twist} = \cos^{-1}(\mathbf{\hat{n_{R_i}}} \cdot \mathbf{\hat{n_{R_j}}}) \quad \text{------------------} \quad \textbf{(9)}$$

Where $\mathbf{\hat{n_{r_i}}}$ and $\mathbf{\hat{n_{r_j}}}$ are unit normal vectors for rings $R_i$ and $R_j$ respectively.

Furthermore, the relative rotation of one ring around the other was quantified by the angle $\phi$ swept by the in-plane vector connecting the centers of mass of the two rings, relative to a fixed reference vector on the reference ring.

**S2. (E) Free Energy Landscape**

The two-dimensional (2D) free energy landscape was reconstructed from the conformational ensemble sampled during the simulations. For a given pair of collective variables, $CV_1$ (twist angle ($\theta_{twist}$) and $CV_2$ (centre-of-mass distance ($d_{COM}$), the joint probability density ρ($CV_1$,

$CV_2$) was estimated using Gaussian kernel density estimation. The corresponding free energy surface was then obtained through Boltzmann inversion:

$$G = -k_B T \ln(\rho) \qquad \text{------------------} \quad \textbf{(10)}$$

$$G_{\text{shifted}} = G - \min(G) \qquad \text{------------------} \quad \textbf{(11)}$$

Where $k_B$ is the Boltzmann constant and T is the simulation temperature. The free energy was shifted such that the global minimum corresponds to zero.

**S3. Evolution of RMSD and $R_g$ during multiscale simulation of the DNA minicircle.**

To characterize conformational deviations and compaction behaviour of the DNA minicircle having different superhelical densities, the RMSD and Rg were monitored throughout the AA and CG simulation trajectories from their initial conformation. In the 0.5 μs AA simulations, the relaxed (σ = 0) and negatively supercoiled (σ = −0.1) minicircles equilibrated to RMSD values of 12–15 Å relative to their initial conformations, whereas the positively supercoiled (σ = +0.1) minicircle exhibited substantially larger deviations (~35 Å) (**Supplementary Figure S3A).** This higher RMSD reflects that the structure is deviating more from the starting structure. Correspondingly, the σ = +0.1 minicircle displayed a lower Rg than the σ = 0 and σ = −0.1 systems **(Supplementary Figure S3B)**, indicating a more compact, writhe-driven conformation arising from positive supercoiling. Over the longer 10 μs CG simulations, both supercoiled topologies (σ = +0.1 and σ = −0.1) showed greater RMSD relative to their initial structures than the relaxed (σ = 0) minicircle, indicating that the introduction of superhelical stress in both supercoil topologies promotes larger-scale deviations. conformational rearrangement on a longer timescale (**Supplementary Figure S3C)**. Among the three, the σ = −0.1 minicircle exhibited the lowest Rg, consistent with enhanced writhing and a correspondingly more compact global conformation relative to the σ = 0 and σ = +0.1 topologies (**Supplementary Figure S3D).**

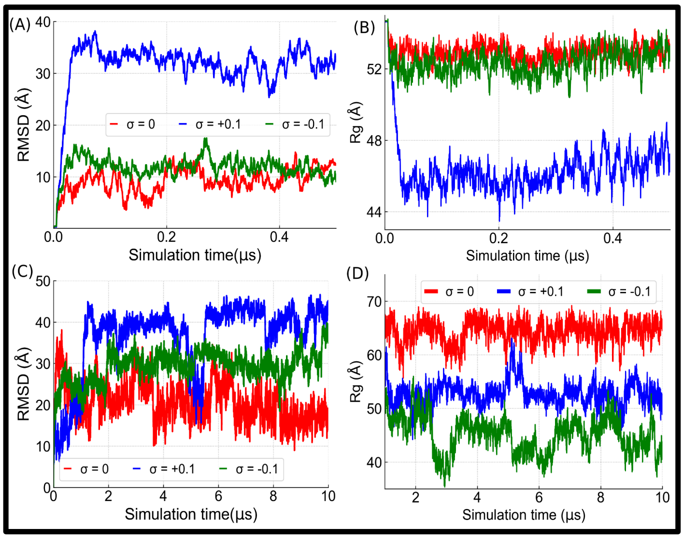


**Supplementary Figure S3. Time evolution of RMSD and Rg for AA and CG simulations of the DNA minicircle at different superhelical densities (σ). (A and B)** RMSD and Rg, respectively, over 0.5 μs of AA simulation for σ = 0 (relaxed), σ = +0.1 (positively supercoiled), and σ = −0.1 (negatively supercoiled) minicircles, calculated relative to their initial conformation. **(C and D)** RMSD and Rg, respectively, over 10 μs of CG simulation for the same three topologies.

### S4. H- bonds in AA simulation of DNA minicircles with varying superhelical densities.

To investigate the effect of supercoiling on duplex stability, the number of canonical Watson–Crick hydrogen bonds (H-bonds) between complementary strands was monitored throughout the 500 ns atomistic simulations **(Supplementary Figure S4).** The relaxed minicircle ($\sigma = 0$) maintained a higher number of H-bonds, with an average of $\langle N_{\mathrm{HB}} \rangle = 191$, indicating preservation of the native B-DNA duplex throughout the simulation. In contrast, both positively ($\sigma = +0.1$) and negatively ($\sigma = -0.1$) supercoiled minicircles exhibited a reduced average H-bond count of approximately 183, accompanied by noticeably larger temporal fluctuations. These observations indicate that torsional stress induces localized destabilization of Watson–Crick base pairing in both overwound and underwound DNA. Although the average number of H-bonds is similar for the two supercoiled systems, the underlying structural mechanisms differ. In the positively supercoiled minicircle, H-bond loss is primarily associated with localized kinks and highly bent regions that relieve excess torsional stress, whereas the negatively supercoiled minicircle exhibits transient denaturation bubbles and local base-pair opening. Such stress-induced structural defects have been reported previously in atomistic simulations of supercoiled DNA minicircles and are recognized as important mechanisms for accommodating torsional strain while preserving the overall duplex architecture (25-27)

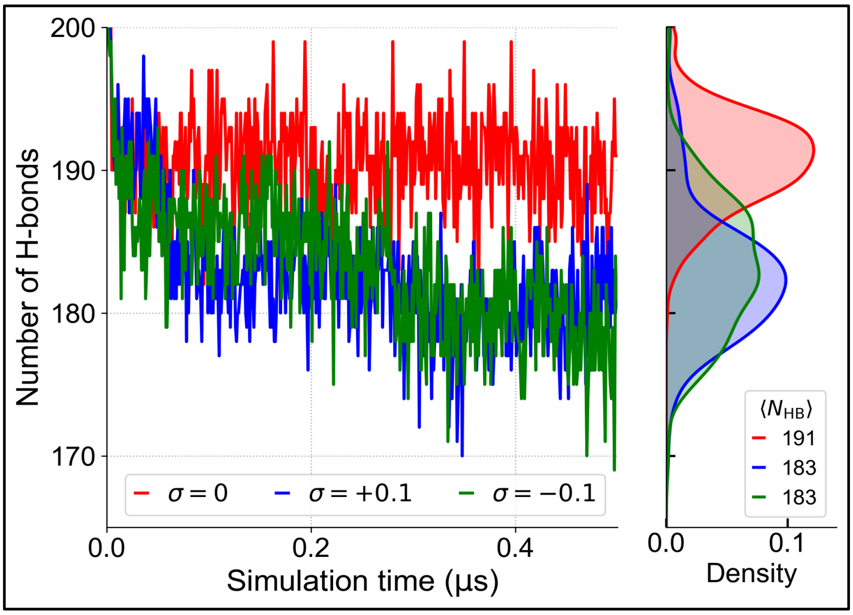


**Supplementary Figure S4.** Time evolution (left) and distribution (right) of the Canonical Watson–Crick hydrogen-bonds between complementary nucleotides in AA simulations of the DNA minicircle at three different superhelical densities.

#### S5. Structural organization of DNA minicircles revealed by contact distance matrix

To characterize the global structural organization of the 100 bp DNA minicircle under different superhelical densities, we computed time-averaged base-pair distance matrices ⟨D⟩ from both all AA and CG simulations (**Supplementary Figure S5).** The scheme for calculating the distance matrix has been explained in the **Supplementary analysis section S2. (A).** In the relaxed DNA (σ = 0), both AA and CG simulations show a smooth, uniform distance distribution dominated by the dark diagonal, reflecting short-range ordering of consecutive base pairs. Introducing positive (σ = +0.1) or negative (σ = −0.1) supercoiling progressively distorts this pattern, with off-diagonal features indicating long-range tertiary contacts and structural compaction driven by torsional stress. Overall, the AA and CG matrices show consistent trends across all superhelical densities, with the CG model additionally capturing larger conformational excursions owing to its extended sampling timescale. In all supercoil densities, the gradual increase in $\langle D_{ij} \rangle$ with sequence separation followed by a decrease at large separations is consistent with the closed circular topology of the minicircle.

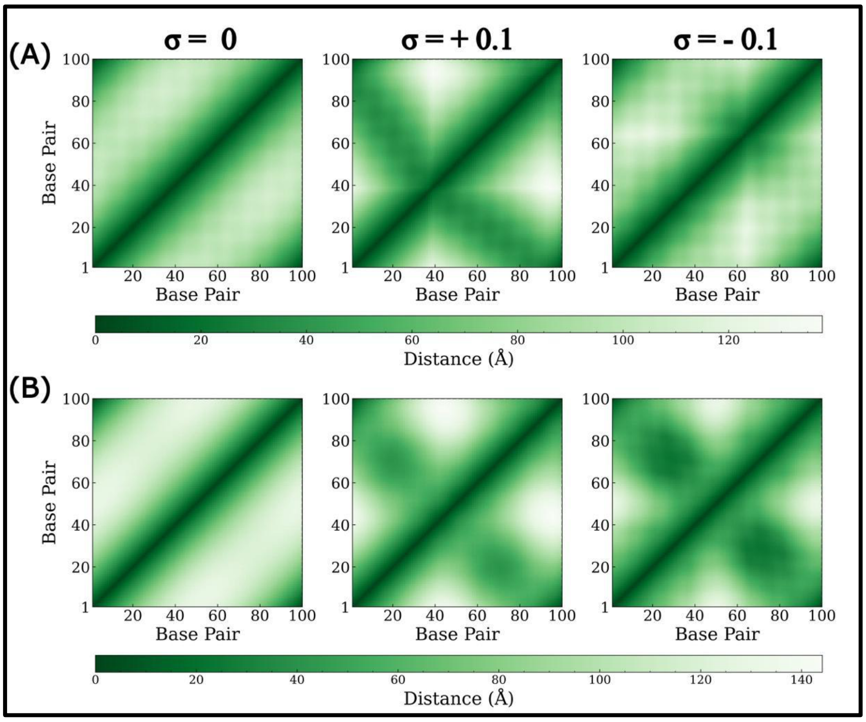


**Supplementary Figure S5. Time-averaged base-pair distance matrices ⟨D⟩ for a 100 bp DNA minicircle obtained from AA and CG simulations under varying superhelical density. (A)** Base-pair distance matrices from AA simulations (0.5 µs) at superhelical densities σ=0 (relaxed), σ= +0.1 (positively supercoiled), and σ= −0.1 (negatively supercoiled). **(B)** Corresponding matrices from CG simulations (10 µs), capturing larger conformational dynamics that were inaccessible at the AA timescale.

#### S6. $Na^+$ ion density distribution in AA simulation around the DNA minicircle

DNA is a highly negatively charged polyelectrolyte due to the presence of phosphate groups along its backbone. This negative charge of the polyanionic backbone is screened by the positively charged salt counterions through electrostatic interaction, leading to neutralization of the backbone's negative charge. While the repulsion of negatively charged salt ions ($Cl^-$). These interactions between the DNA backbone and counterions can be either directly as chelated ions (28) or through shells of coordinated water molecules called diffuse ions or "ion atmosphere" (29). Despite the diversity of ions capable of interacting with DNA in biophysical systems, $K^+$, $Na^+$, $Mg^{2+}$, and $Cl^-$ are among the most physiologically relevant ions and are therefore commonly employed in molecular dynamics simulations. In this study, $Na^+$ and $Cl^-$

ions were used to neutralize the system and maintain the desired salt concentration as routinely used in the experiments.

In order to obtain an insight into how $Na^+$ ions are spatially organized around the minicircle over the course of the simulation, we computed time-averaged three-dimensional ion density maps projected onto two different planes XY, XZ, and ZY, **Supplementary Figure S6**.

Across all superhelical states, the $Na^+$ ion density remains tightly localized near the DNA backbone, consistent with Manning counterion condensation theory (30,31) and shielding the negative charge of DNA backbone. Details of the ion density calculation are provided in **Supplementary Section S2(B)**.

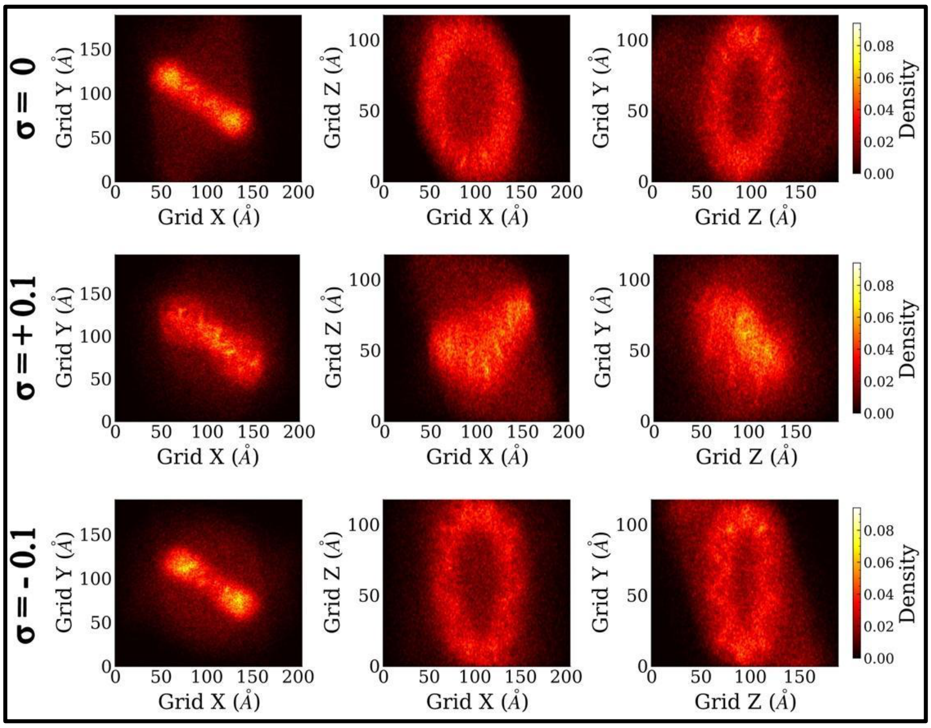


**Supplementary Figure S6. $Na^+$ ion density distribution around DNA minicircle obtained from AA simulations under varying superhelical density.** Two-dimensional (2D) projections of the three-dimensional $Na^+$ ion density field accumulated around the DNA onto two planes XY, XZ, and ZY are shown for three superhelical densities: relaxed (σ = 0, top row), positively supercoiled (σ = +0.1, middle row), and negatively supercoiled (σ = −0.1, bottom row). The color scale encodes the time-averaged $Na^+$ ion density around the DNA accumulated over the 0.5 µs trajectory.

### S7. $Na^+$ ion distribution in CG simulations around the DNA minicircle

Compared with the AA simulations, the Martini2 CG model exhibits a substantially broader $Na^+$ ion atmosphere around the DNA across all superhelical states **Supplementary Figure S7.** Although $Na^+$ ions are represented as singly charged beads, the coarse-grained representation of DNA and the reduced electrostatic resolution of the Martini model result in a more spatially averaged ion atmosphere around the DNA. Furthermore, the relatively large bead size limits the ability of ions to access DNA grooves and reproduce the detailed ion-phosphate interactions observed in atomistic models. The original Martini force-field paper by **Uusitalo *et al., 2015 (15)*** noted that the Martini DNA model successfully captures many structural properties of DNA; the coarse-grained treatment inherently reduces the level of detail in ion-mediated interactions. Consistent with this observation, **Naskar *et al.* (2021)** (32) reported that coarse-grained DNA models do not fully reproduce salt-dependent DNA properties due to limitations in describing detailed electrostatic and ion-correlation effects. These factors likely contribute to the diffuse $Na^+$ density distributions observed in the Martini2 simulations relative to the all-atom model.

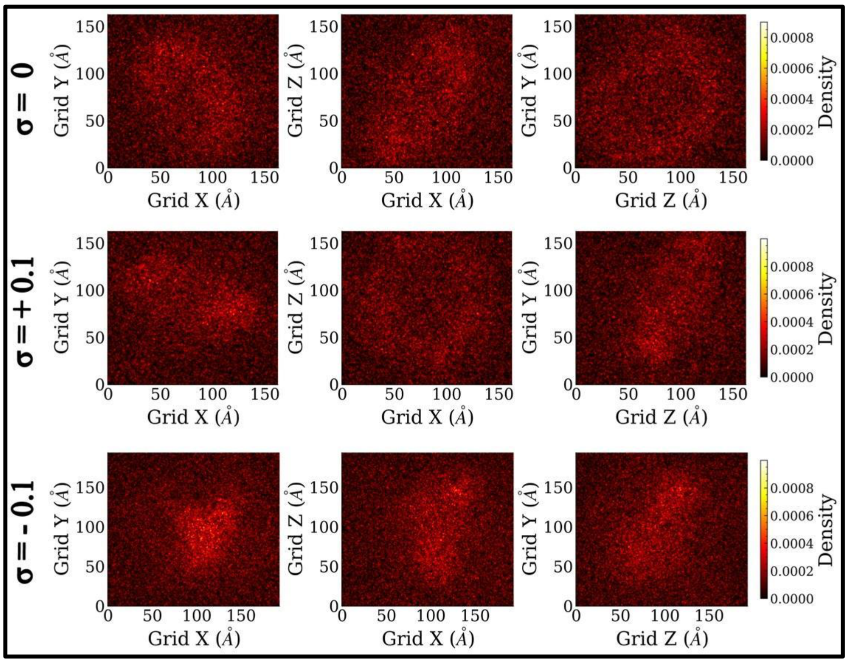


**Supplementary Figure S7. $Na^+$ ion density distribution around DNA minicircle obtained from CG simulations under varying superhelical density.** 2D projections of the three-dimensional $Na^+$ ion density accumulated around the DNA onto two planes, XY, XZ, and ZY are shown for three superhelical densities: relaxed ($\sigma$ = 0, top row), positively supercoiled ($\sigma$ = +0.1, middle row), and negatively supercoiled ($\sigma$ = −0.1, bottom row). The color scale encodes the time-averaged $Na^+$ ion density around the DNA accumulated over the 10 μs trajectory.

**S8. Time evolution of the RMSD and $R_g$ during AA simulation of DNA catenanes.**

To assess the structural stability of the individual DNA rings, chain-wise RMSD and Rg were monitored throughout the AA simulations **(Supplementary Figure S8)**. For the homocatenane, both 100 bp rings exhibit an increase in RMSD. Ring 1 has a higher RMSD than ring 2, as it deviates more from the initial reference conformation. RMSD also reflects that both rings are showing different dynamics **(Supplementary Figure S8A)**. To measure compactness, the $R_g$ values for both rings were quantified, and it remained nearly constant at approximately 53 Å throughout the simulation (**Supplementary Figure S8B**). The distribution of $R_g$ and its full width at half maximum (FWHM) also shows little difference.

For the heterocatenane, the 200 bp ring 1 displays higher RMSD deviations than the 100 bp ring 2 (**Supplementary Figure S8C**), consistent with the greater flexibility expected for the longer DNA contour length. The $R_g$ analysis also further supports this trend that ring 1 with more bps has more $R_g$ as compared to ring 2 with fewer bps, and it explores more conformations (**Supplementary Figure S8D**). Ring 1 shows more distribution, and its FWHM is 1.98 Å, while ring 2 shows a narrow distribution and its FWHM is 0.80 Å.

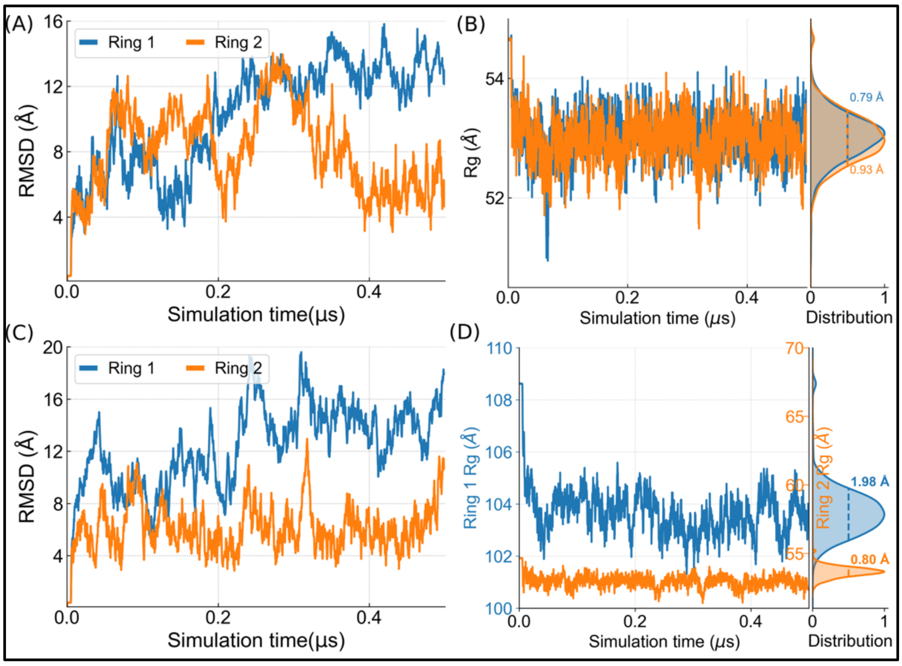


**Supplementary Figure S8. Time evolution of RMSD and Rg of the homo- and hetero catenanes in AA simulations. (A, C)** Ring-wise RMSD profile of the homo (100 + 100 bp) and hetero catenane (200 + 100 bp), respectively, as a function of simulation time, calculated relative to their initial conformations. **(B, D)** Ring-wise Rg time series and their corresponding distribution for the homo- and heterocatenanes, respectively. In the distribution plots, the full width at half maximum (FWHM) for each ring is shown as a dashed line.

### S9. Topological fluctuations in DNA homocatenane during multiscale simulations

The evolution of writhe (ΔWr) and twist (ΔTw) as a function of simulation time was quantified in AA and CG simulations to characterize the global topology and torsional state of the DNA rings during the simulations **(Supplementary Figure S9)**. As the linking number of closed DNA remains conserved, changes in twist are compensated by changes in writhe, providing insight into the redistribution of superhelical stress and large-scale conformational dynamics while preserving the overall topology.

In the AA simulations, both rings exhibit only small positively writhe fluctuations throughout the trajectory **(Supplementary Figure S9A)**, indicating that the homocatenane remains largely planar and tries to writhe positively without undergoing significant global buckling or supercoiling. In the CG simulation, both rings are writhing negatively (**Supplementary Figure S9B)**. Correspondingly, the evolution of the change in writhe (ΔWr) and twist (ΔTw) for rings 1 (left) and 2 (right) during the AA simulations demonstrates that minor fluctuations in ΔTw is compensated by a corresponding positive increase in ΔWr, reflecting the dynamic redistribution of torsional stress in the DNA rings **(Supplementary Figure S9C).**

The CG simulations show larger-amplitude fluctuations in both ΔTw and ΔWr than the AA simulations, indicating enhanced sampling of global topological conformations over the extended 10 μs timescale **(Supplementary Figure 9D)**. Both the rings are showing strong negative writhing throughout the entire simulation time to leads to formation of plectonemic-like structures in the CG simulation **(Supplementary movie SM2 )** This compensatory interplay between twist and writhe is a fundamental feature of closed DNA topology and is also observed in vivo, where DNA supercoiling is continuously regulated by DNA-binding proteins and topoisomerases (33).

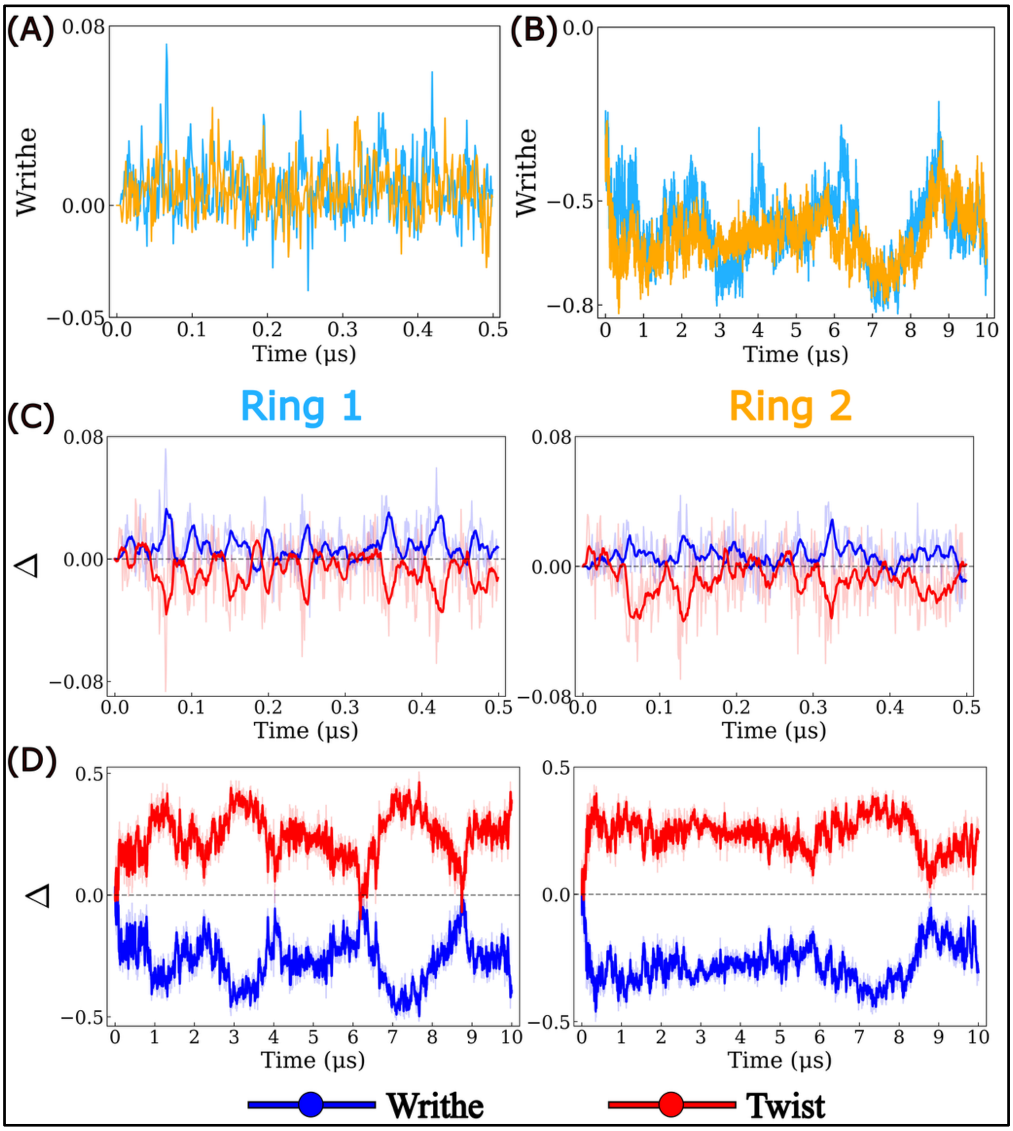


**Supplementary Figure S9: Evolution of DNA writhe and the coupled changes in writhe (ΔWr) and twist (ΔTw) for the homocatenane during AA and CG simulations (see also the supplementary analysis section S2C). (A-B)** Time evolution of writhe for rings 1 (cyan) and 2 (orange) from the AA (0.5 μs) and CG (10 μs) simulations, respectively. **(C-D)** Time evolution of the changes in writhe (ΔWr, blue) and twist (ΔTw, red) for rings 1 (left) and 2 (right) from the AA and CG simulations, respectively.

### S10. Topological fluctuations DNA heterocatenane during multiscale simulations

The topological dynamics of the two rings in the DNA heterocatenane are characterized by the evolution of writhe (Wr), the change in writhe (ΔWr), and the twist (ΔTw) **(Supplementary Figure S10)**.  During the 0.5 μs AA simulation, both rings exhibit relatively small writhe fluctuations around their initial values**.** Ring 1 samples both positive and negative writhe values, whereas Ring 2 remains close to zero with a slight positive bias. It indicates that the rings largely preserve their initial conformations and undergo only local topological rearrangements **(Supplementary Figure S10A)**. In contrast, the 10 μs CG simulation captures enhanced topological dynamics, with Ring 1 predominantly adopting positive writhe and Ring 2 negative writhe, reflecting greater conformational sampling over longer timescales **(Supplementary Figure S10B).**

The changes in writhe (ΔWr) and twist (ΔTw) further demonstrate the coupling between these two topological parameters. During the AA simulation, Ring 1 exhibits continuous exchange between writhe and twist, while Ring 2 shows weaker but similar compensatory fluctuations **(Supplementary Figure S10C)**. A comparable trend is observed in the CG simulation for ring 1, although the magnitude of the fluctuations is substantially larger **(Supplementary Figure S10D).** Ring 1 continues to show dynamic interconversion between the writhe and twist. While Ring 2 displays a stable increase in positive twist, accompanied by negative writhe.

Overall, the multiscale simulations demonstrate that changes in twist are compensated by opposing changes in writhe, illustrating the redistribution of topological stress. The corresponding dynamics are shown in **Supplementary Movie 2 (SM2)**.

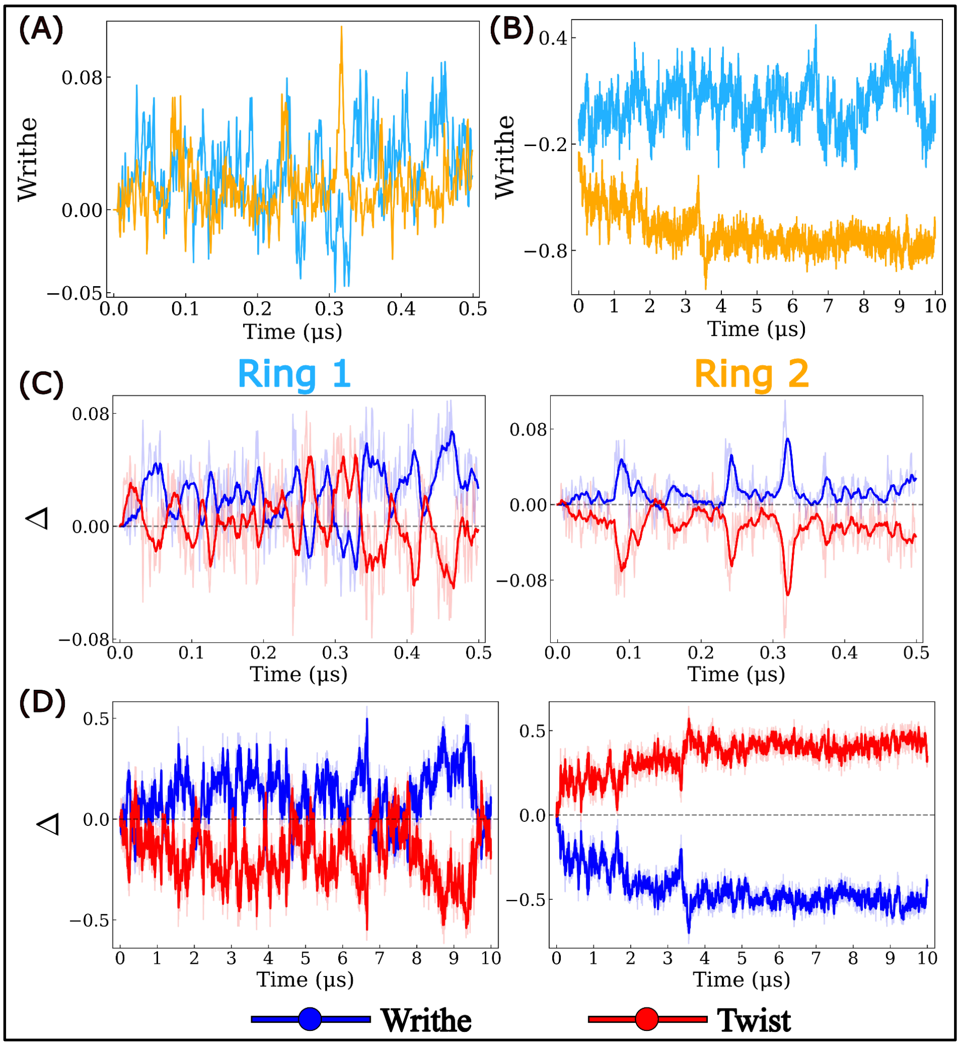


**Supplementary Figure S10. Evolution of DNA writhe and the coupled changes in writhe (ΔWr) and twist (ΔTw) for the heterocatenane during AA and CG simulations (see also the supplementary analysis section S2C). (A-B)** Time evolution of writhe for rings 1 (cyan) and 2 (orange) from the AA (0.5 μs) and CG (10 μs) simulations, respectively. **(C-D)** Time evolution of the changes in writhe (ΔWr, blue) and twist (ΔTw, red) for rings 1 (left) and 2 (right) from the AA and CG simulations, respectively.

#### S11. 2D Free Energy of DNA catenanes deduced from multiscale simulations

The 2D Free Energy Landscapes (FEL) from the AA simulations **(Supplementary Figure S11A and C)** shows how the system evolves from an initial configuration in which the two rings are nearly perpendicular (black dot). The global free energy minimum (green circle) lies close to the initial conformation (black circle) for both the homo- and heterocatenanes, indicating that only modest changes in the twist angle and $d_{COM}$ occur during the 0.5 µs simulations. The well-defined low-energy basins suggest that both systems remain structurally stable while sampling a limited conformational space.

In contrast, the FEL from the CG simulations **(Supplementary Figure S11B and D)** exhibit broader low-energy basins, reflecting enhanced conformational sampling over the 10 µs simulations. For both catenanes, the global free energy minimum is shifted to higher twist angles and lower $d_{COM}$ values relative to the initial conformation, indicating that the systems preferentially adopt more compact and twisted low-energy states. These observations are consistent with the ΔWr and ΔTw analyses, which show increased writhing and conformational flexibility in the CG simulation.

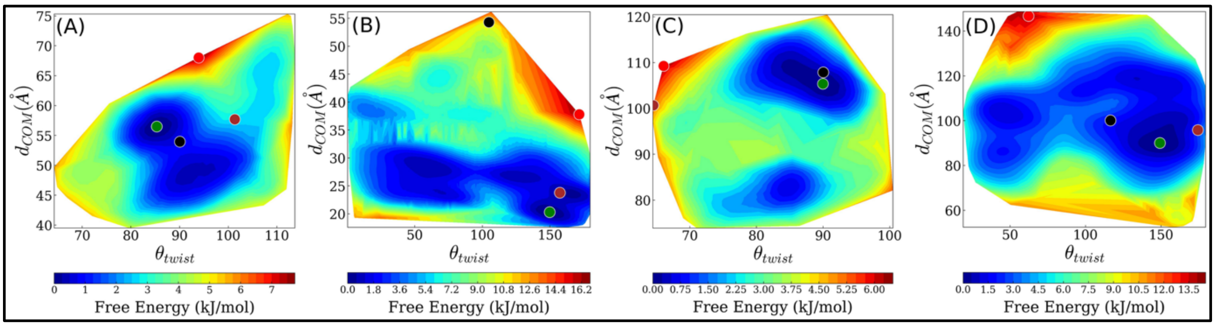


**Supplementary Figure S11. Two-dimensional free-energy surface of DNA catenanes projected onto the centre-of-mass distance ($d_{COM}$) between the rings and the relative twist angle ($\theta_{twist}$) using the Boltzmann inversion method. (A-B)** FELs for homocatenanes from AA and CG simulations, respectively. **(C-D)** FELs for hetero-catenane from AA and CG simulations, respectively. The black circle denotes the initial conformation, the brown circle marks the conformation at the end of the simulation, the green circle indicates the global free energy minimum, and the red circle represents the highest free energy state. The colored contours represent the free energy (kJ/mol).

### S12. $Na^+$ ion distribution around DNA homocatenane from AA and CG MD simulations

In the homocatenane system, the AA simulations reveal a pronounced $Na^+$ ion atmosphere that closely follows the topology of the interlocked DNA rings, with enhanced ion accumulation in regions where the two negatively charged backbones approach each other , **Supplementary Figure S12 (A)**. In contrast, the Martini2 coarse-grained simulations exhibit a markedly altered ion distribution characterized by sparse, localized density hotspots and the absence of a continuous counterion cloud surrounding the DNA, **Supplementary Figure S12 (B)**. This difference likely arises from the reduced electrostatic resolution of the coarse-grained representation, which limits the ability of the model to capture the detailed ion-mediated correlations and counterion condensation observed in atomistic simulations (15,31).

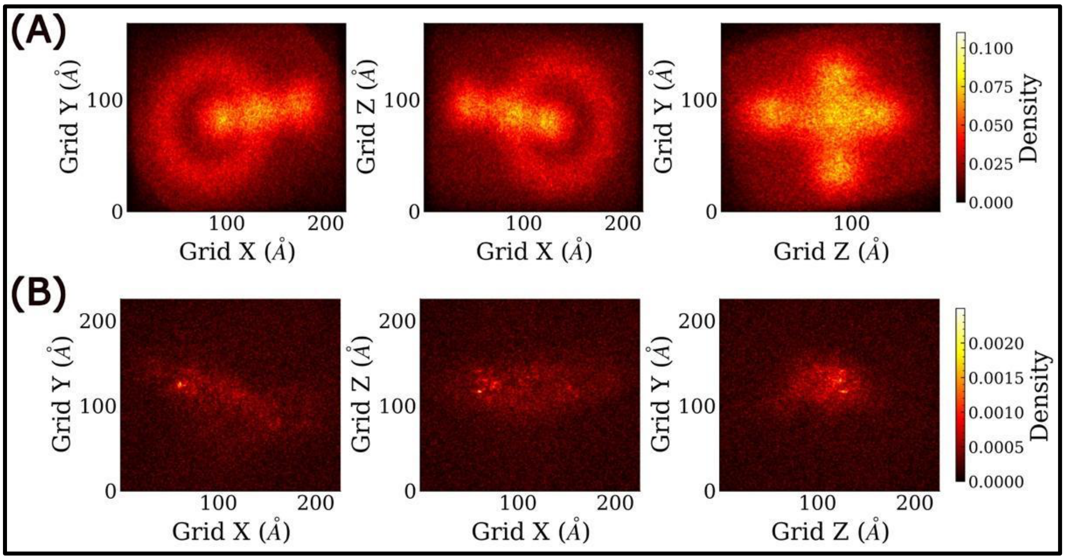


**Supplementary Figure S12. Density of $Na^+$ ion around DNA homocatenane.** Two-dimensional projections of the three-dimensional $Na^+$ ion density field accumulated around the DNA homocatenane (100 bp) onto the XY, XZ, and ZY planes, from **(A)** AA and **(B)** CG simulations, respectively. The color scale encodes the time-averaged $Na^+$ ion number density around the homocatenane accumulated over the 0.5 µs (AA) and 10 µs (CG) trajectory.

### S13. $Na^+$ ion distribution around DNA heterocatenane from AA and CG MD simulations

Consistent with the homocatenane, the heterocatenane also displays a similar ion distribution pattern **(Supplementary Figure S13)**. The AA simulations reveal a pronounced $Na^+$ ion atmosphere surrounding the DNA rings, whereas the Martini2 CG simulations exhibit altered ion distributions characterized by a less continuous and more localized counterion density around the DNA **(Supplementary Figure S13 A,B).**

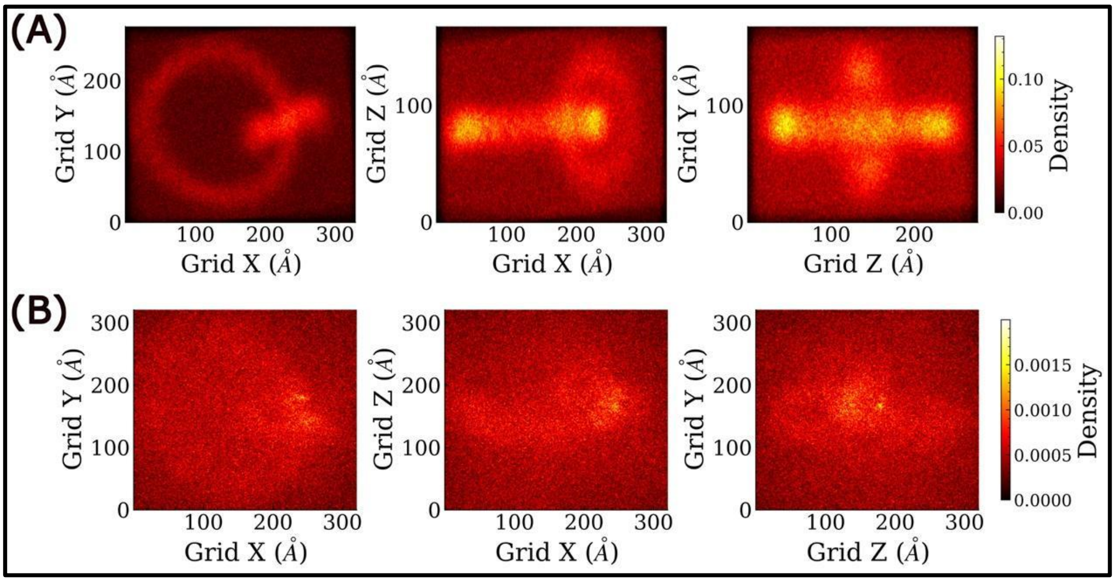


**Supplementary Figure S13. Spatial distribution of $Na^+$ ion around DNA hetero (200 and 100 bps) catenane from AA and CG simulations. (A and B)** Two-dimensional projections of the three-dimensional $Na^+$ ion density field accumulated around the DNA catenane rings onto two planes XY, XZ, and ZY, from AA and CC simulations, respectively. The color scale encodes the time-averaged $Na^+$ ion number density around the hetero catenane rings accumulated over the 0.5 µs (AA) and 10 µs (CG) trajectory.

### S14. Evolution of the RMSD and Rg during CG simulation of DNA catenane

The ring-wise RMSD and Rg of homo- and heterocatenane were calculated throughout the 10 µs CG simulations **(Supplementary Figure S14)**. For the homocatenane, both 100 bp rings undergo an initial structural relaxation followed by RMSD fluctuations, Ring 2 exhibiting slightly larger deviations than Ring 1 (**Supplementary Figure S14 A**). The evolution of Rg values shows that ring 2 is having lesser Rg as compared to ring 1, we can see this clearly through their distribution also. The FWHM of ring 1 is 4.11 Å while for ring 2 it is 3.08 Å. Despite these conformational rearrangements, both rings maintain comparable Rg values throughout the simulation (**Supplementary Figure S14 B**), indicating that their overall compactness is largely preserved.

In the heterocatenane, the 200 bp ring exhibits substantially larger RMSD fluctuations than the 100 bp ring, reflecting its greater conformational flexibility and ability to sample diverse global conformations during the 10 μs trajectory (**Supplementary Figure S14 C**). Consistent with these structural rearrangements, the Rg of the 200 bp ring displays larger fluctuations than that of the 100 bp ring (**Supplementary Figure S14 D**), whereas the smaller ring maintains an approximately constant Rg throughout the simulation. These results indicate that the extended timescale accessible to the CG model enhances conformational sampling, particularly for the larger DNA ring.

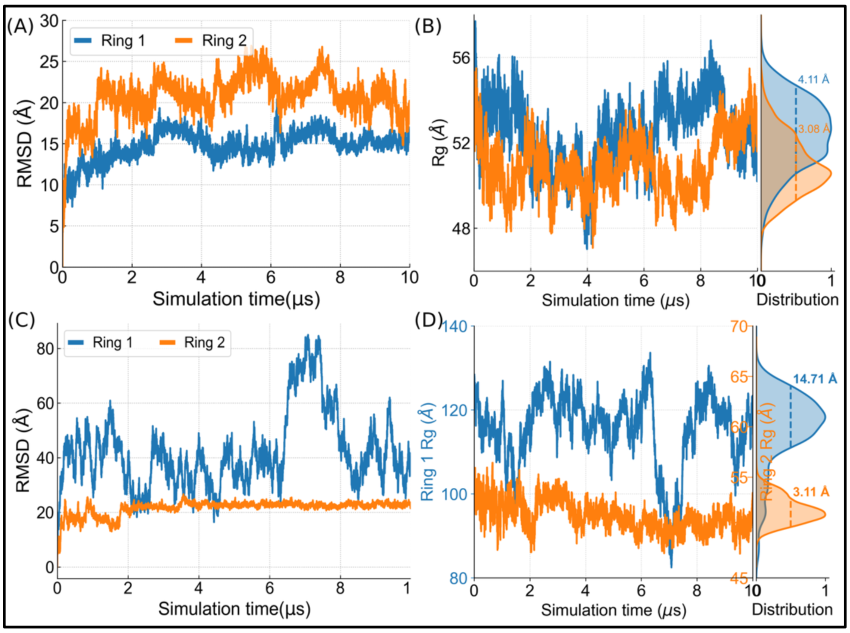


**Supplementary Figure S14. Time evolution of RMSD and Rg during CG simulations of the homo- and hetero catenanes. (A, C)** Ring-wise RMSD profile of the homo- (100 + 100 bp) and hetero catenane (200 + 100 bp), respectively, as a function of simulation time, calculated relative to their initial conformations. **(B, D)** Ring-wise Rg time series and their corresponding distribution for the homo- and hetero-catenanes, respectively. In the distribution plots, the full width at half maximum (FWHM) for each ring is shown as a dashed line.

#### S15. Ring-wise RMSD and $R_g$ analyses of the DNA Borromean rings.

To characterize the conformational stability and compactness of the three DNA rings in Borromean, the ring-wise RMSD and $R_g$ of were calculated throughout the AA and CG trajectories **(Supplementary Figure S15).** During the AA simulations **(Supplementary Figure S15 A-B**), each 120 bp ring underwent an initial relaxation phase followed by an increase in RMSD and kept on fluctuating with 0.5 μs simulation time. While Ring 2 displayed marginally higher deviations compared to other two rings, the $R_g$ for all three rings remained nearly constant, with ring 2 having slightly lower Rg as compared to others, suggesting that the global compactness of the individual components is well-preserved despite localized structural fluctuations. ring-wise RMSD indicating all three rings are showing different dynamics,

Over the 10 μs CG simulations **(Supplementary Figure S15 C-D)**, the rings explored a more diverse conformational ensemble, resulting in larger RMSD values than those observed at the atomistic level. After the equilibration period, the RMSD for each ring remained stable with moderate temporal fluctuation. All three rings are showing different $R_g$ and compacting differently from the initial conformation. The CG model captures broader conformational dynamics over extended timescales.

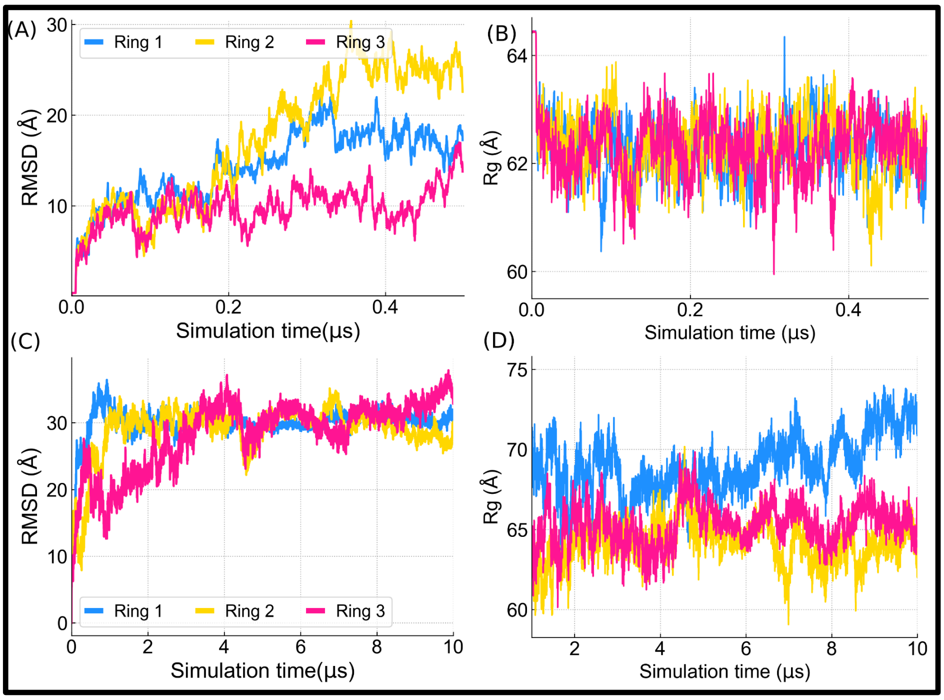


**Supplementary Figure S15.** Time evolution of the RMSD and $R_g$ during AA and CG simulations of the Borromean rings. **(A, C)** Ring-wise RMSD profiles of the three 120-bp rings from the AA and CG simulations, respectively, calculated relative to their initial conformations. **(B, D)** Ring-wise Rg profiles of the three rings from the AA and CG simulations, respectively, shown as a function of simulation time.

### S16. Structural organization of Borromean rings revealed by contact distance matrix

The time-averaged base-pair distance matrices ⟨D⟩, **Supplementary Figure S16** shows the AA matrix displays smooth intra- and inter-ring distance gradients consistent, while the CG matrix exhibits a pronounced checkerboard pattern of alternating contact and separation regions, reflecting large-scale correlated structural rearrangements between the three rings over the longer simulation timescale. Overall, CG model additionally captures larger conformational excursions owing to its extended sampling timescale.

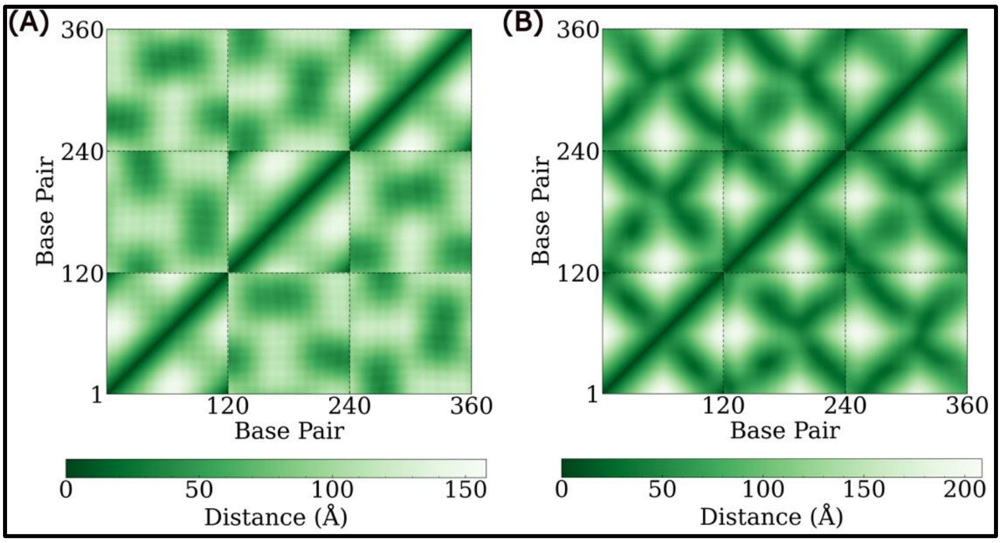


**Supplementary Figure S16.** Time-averaged base-pair distance matrices ⟨D⟩ for a 120 bp DNA Borromean rings obtained from AA and CG simulations.

**S17. Evolution of writhe and twist in individual rings of the Borromean DNA assembly**

To investigate the topological dynamics of the Borromean DNA assembly, we monitored the writhe (Wr) of each ring together with the corresponding changes in twist (ΔTw) and writhe (ΔWr) throughout the AA and CG simulations. During the 0.5 μs AA simulations, the three rings exhibited only modest fluctuations in writhe, reflecting the limited conformational sampling accessible at the atomistic timescale **(Supplementary Figure S17 A)**. The change in twist ( ΔTw) and writhe (ΔWr) in all three rings are very less. The average topological parameters further show that the rings accommodate torsional stress primarily through a gradual increase in writhe, resulting in a positively supercoiled state throughout the simulation **(Supplementary Figure S17C)**. Although all three rings maintained similar positive writhe values, no significant plectoneme formation was observed.

In contrast, the 10 μs CG simulations sampled a substantially larger conformational space, leading to pronounced and persistent changes in writhe for all three rings **(Supplementary Figure S17 B)**. The change in twist ( ΔTw) and writhe (ΔWr) from the CG simulation shows

**(Supplementary Figure S17 D)** ring 1 initially writhe positively, which subsequently relaxed and fluctuated around a small positive value throughout the remainder of the simulation. Ring 2 exhibited the largest topological rearrangement, maintaining strongly negative writhe that was compensated by a corresponding increase in twist, indicative of sustained twist–writhe interconversion. In comparison, Ring 3 displayed more moderate fluctuations, with transient changes in writhe and twist while remaining centered near zero. The development of negative supercoiling promoted plectoneme formation, causing the rings to evolve from a mutually orthogonal arrangement to a more tilted configuration with persistent inter- and intra-ring contacts.

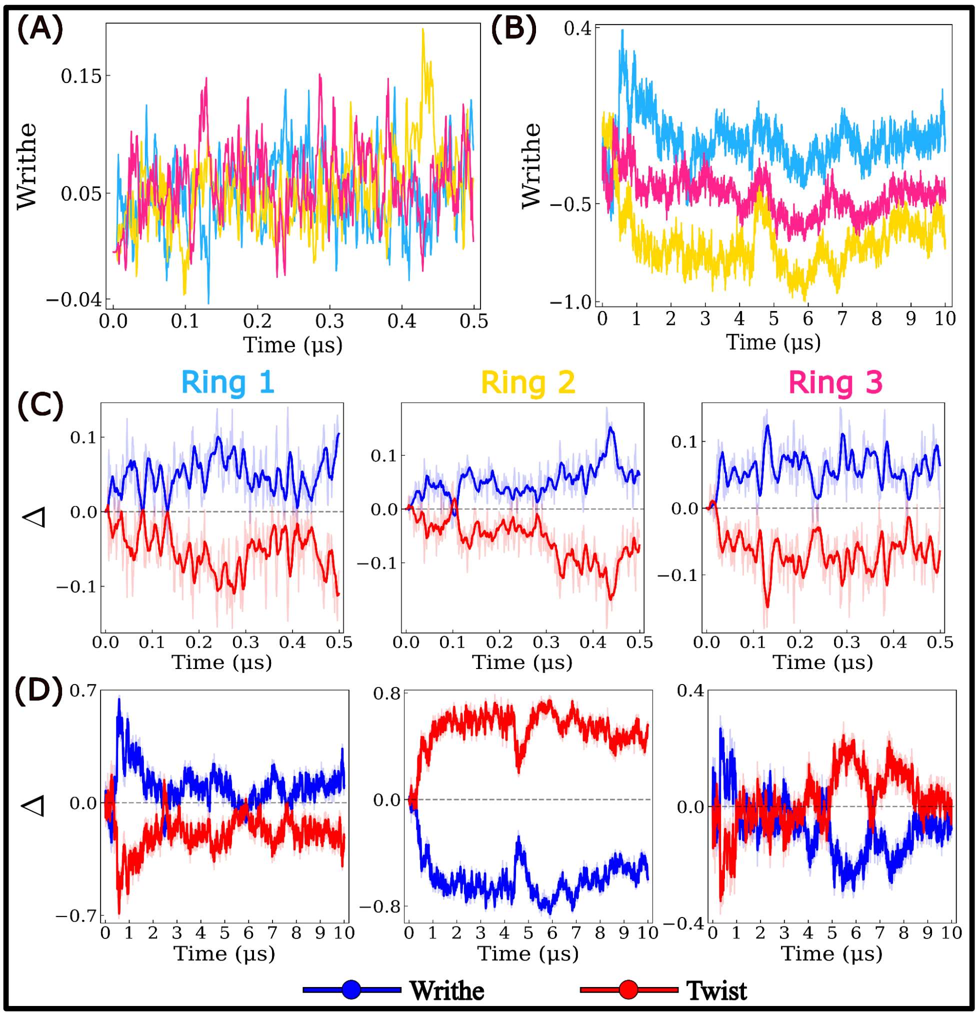


**Supplementary Figure S17. Time evolution of writhe and twist-writhe changes in borromean rings from AA and CG simulations. (A and B)** Evolution of writhe in all three individual rings from AA and CG simulations, respectively. **(C and D)** Change in twist (ΔTw) and writhe (ΔWr) with simulation time in individual all three rings, from AA and CG simulations, respectively. Here ΔTw is shown in red, while

ΔWr in blue color. shaded lines represent raw data while solid lines represent smoothed trajectories. The dashed horizontal line at Δ=0 serves as reference for the initial state.

#### S18. Supplementary Movies:

**Dropbox links for the video tutorial (SM1) and simulation movies (SM2-SM4)**

[https://www.dropbox.com/scl/fo/vyz7sn82j5hcp6vfpzoyg/AGscE_p-3zQpVsAEOaUhNUY?rlkey=wjjxaeolu1tpc4biesna1vrds&st=eharwmq8&dl=0](https://www.dropbox.com/scl/fo/vyz7sn82j5hcp6vfpzoyg/AGscE_p-3zQpVsAEOaUhNUY?rlkey=wjjxaeolu1tpc4biesna1vrds&st=eharwmq8&dl=0)

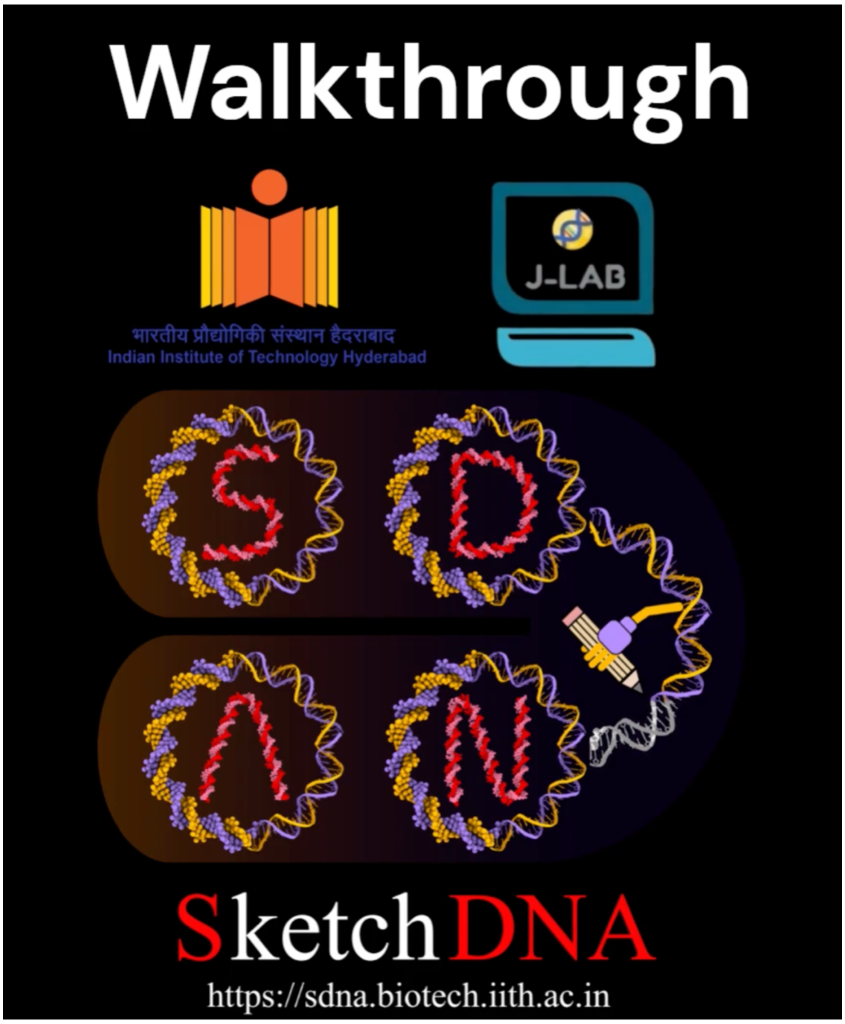


**Supplementary Movie 1 (SM1) –** This video tutorial provides a quick tour of the interface and basic options available in the SDNA toolkit.

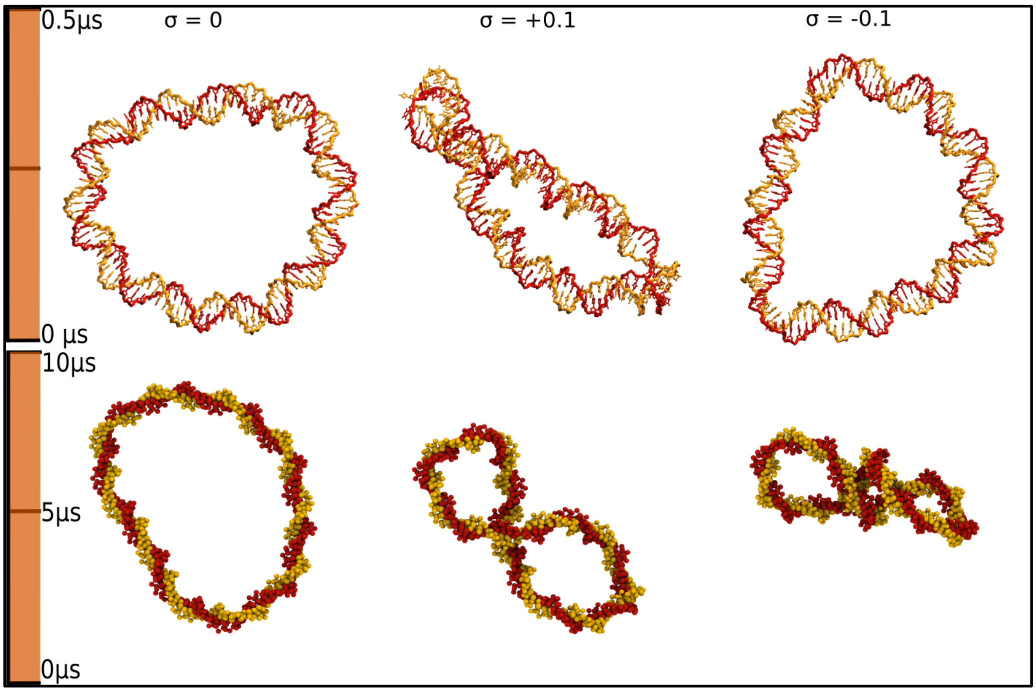


**Supplementary Movie 2 (SM2). Supercoiling-dependent structural dynamics of 100 bp DNA minicircles in all-atom (AA) and coarse-grained (CG) simulations:** A movie illustrating the trajectories of 100 bp DNA minicircles with different superhelical densities: relaxed (σ = 0), positively supercoiled (σ = +0.1), and negatively supercoiled (σ = −0.1), obtained from AA (top) and CG (bottom) molecular dynamics simulations. AA simulations were performed for 0.5 µs, whereas CG simulations were performed for 10 µs for each system. The snapshots shown correspond to the final configurations of the trajectories, illustrating the distinct conformational responses of DNA minicircles to different supercoiling states. The two DNA strands are colored golden yellow and red. DNA is represented in licorice and van der Waals (VdW) representations for the AA and CG models, respectively. Water molecules and ions are omitted for clarity.

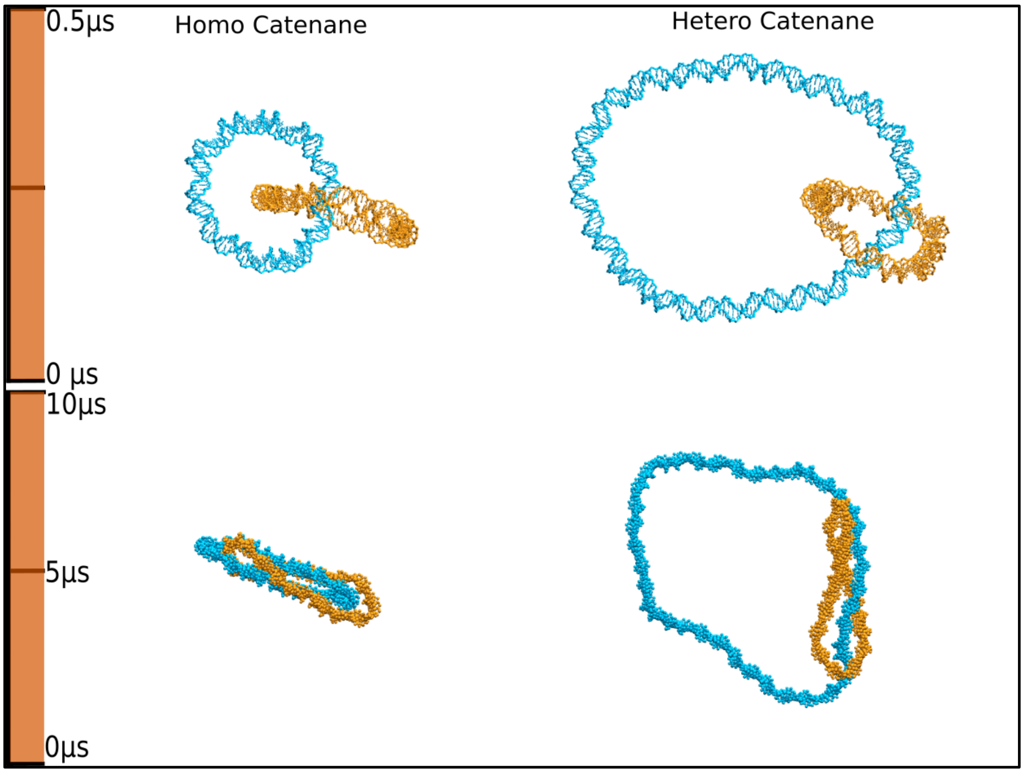


**Supplementary Movie 3 (SM3). Trajectories of homo- and hetero-DNA catenanes obtained from AA and CG MD simulations:** The homocatenane consists of two interlocked 100 bp DNA rings, whereas the hetero-catenane comprises a 200 bp DNA ring interlocked with a 100 bp DNA ring. All rings have zero superhelical density (σ = 0). The DNA rings are colored cyan and orange to distinguish the two rings within each catenane. The top panels show AA trajectories (0.5 µs), while the bottom panels show CG trajectories (10 µs). The movies illustrate the conformational fluctuations and relative motions of the interlocked rings while preserving the catenated topology. DNA is represented in licorice (AA) and VdW (CG) representations. Water molecules and ions are omitted for clarity.

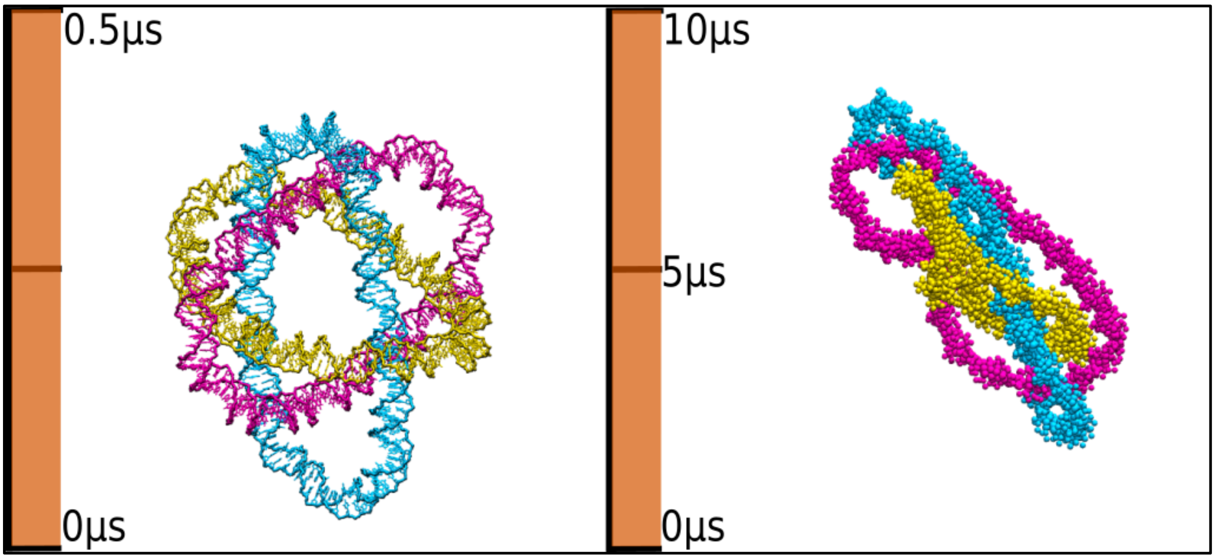


**Supplementary Movie 4 (SM4). Trajectories of a DNA borromean ring obtained from AA and CG MD simulations:** The Borromean ring consists of three interlocked 120 bp DNA rings with zero superhelical density (σ = 0). Although no two rings are directly linked, the three-ring assembly remains topologically linked as a whole. The left panel shows the AA trajectories (0.5 μs), while the right panel shows the CG trajectories (10 μs). The movie illustrates the conformational fluctuations and collective motions of the three-ring assembly while preserving the borromean topology. The three DNA rings are shown in cyan, magenta, and yellow. DNA is represented in licorice (AA) and VdW (CG) representations. Water molecules and ions are omitted for clarity.

### S19. References


1. Case, D.A., Aktulga, H.M., Belfon, K., Cerutti, D.S., Cisneros, G.A., Cruzeiro, V.W.D., Forouzesh, N., Giese, T.J., Götz, A.W. and Gohlke, H. (2023) AmberTools. *Journal of chemical information and modeling*, **63**, 6183-6191.
2. Case, D.A., Cheatham III, T.E., Darden, T., Gohlke, H., Luo, R., Merz Jr, K.M., Onufriev, A., Simmerling, C., Wang, B. and Woods, R.J. (2005) The Amber biomolecular simulation programs. *Journal of computational chemistry*, **26**, 1668-1688.
3. Darden, T., York, D. and Pedersen, L. (1993) Particle mesh ewald - An N.log(N) method for ewald sums in large systems. *Journal of Chemical Physics*, **98**, 10089-10092.

4. Zgarbová, M., Sponer, J. and Jurecka, P. (2021) Z-DNA as a touchstone for additive empirical force fields and a refinement of the alpha/gamma DNA torsions for AMBER. *Journal of Chemical Theory and Computation*, **17**, 6292-6301.
5. Jorgensen, W.L., Chandrasekhar, J., Madura, J.D., Impey, R.W. and Klein, M.L. (1983) Comparison of simple potential functions for simulating liquid water. *The Journal of chemical physics*, **79**, 926-935.
6. Joung, I.S. and Cheatham III, T.E. (2008) Determination of alkali and halide monovalent ion parameters for use in explicitly solvated biomolecular simulations. *The journal of physical chemistry B*, **112**, 9020-9041.
7. Yoo, J. and Aksimentiev, A. (2012) Improved parametrization of Li+, Na+, K+, and Mg2+ ions for all-atom molecular dynamics simulations of nucleic acid systems. *The journal of physical chemistry letters*, **3**, 45-50.
8. Davidchack, R.L., Handel, R. and Tretyakov, M. (2009) Langevin thermostat for rigid body dynamics. *The Journal of chemical physics*, **130**.
9. Miyamoto, S. and Kollman, P.A. (1992) Settle: An analytical version of the SHAKE and RATTLE algorithm for rigid water models. *Journal of computational chemistry*, **13**, 952-962.
10. Andersen, H.C. (1983) Rattle: A “velocity” version of the shake algorithm for molecular dynamics calculations. *Journal of Computational Physics*, **52**, 24-34.
11. Salomon-Ferrer, R., Gotz, A.W., Poole, D., Le Grand, S. and Walker, R.C. (2013) Routine microsecond molecular dynamics simulations with AMBER on GPUs. 2. Explicit solvent particle mesh Ewald. *Journal of chemical theory and computation*, **9**, 3878-3888.
12. Humphrey, W., Dalke, A. and Schulten, K. (1996) VMD: Visual molecular dynamics. *J. Mol. Graph.*, **14**, 33-38.
13. Pettersen, E.F., Goddard, T.D., Huang, C.C., Meng, E.C., Couch, G.S., Croll, T.I., Morris, J.H., and Ferrin, T.E. (2021) UCSF ChimeraX : Structure visualization for researchers, educators, and developers. *Protein Science*, **30(1)**, 70-82.
14. Roe, D.R. and Cheatham, T.E., III. (2013) PTRAJ and CPPTRAJ: Software for Processing and Analysis of Molecular Dynamics Trajectory Data. *Journal of Chemical Theory and Computation*, **9**, 3084-3095.
15. Uusitalo, J.J., Ingólfsson, H.I., Akhshi, P., Tieleman, D.P. and Marrink, S.J. (2015) Martini coarse-grained force field: extension to DNA. *Journal of chemical theory and computation*, **11**, 3932-3945.

16. Marrink, S.J., Risselada, H.J., Yefimov, S., Tieleman, D.P. and De Vries, A.H. (2007) The MARTINI force field: coarse grained model for biomolecular simulations. *The journal of physical chemistry B*, **111**, 7812-7824.
17. Abraham, M.J., Murtola, T., Schulz, R., Páll, S., Smith, J.C., Hess, B. and Lindahl, E. (2015) GROMACS: High performance molecular simulations through multi-level parallelism from laptops to supercomputers. *SoftwareX*, **1**, 19-25.
18. Abraham, M., Alekseenko, A., Bergh, C., Blau, C., Briand, E., Doijade, M., Fleischmann, S., Gapsys, V., Garg, G. and Gorelov, S. (2023) GROMACS 2023.3 Source code. *Zenodo*.
19. Bussi, G., Donadio, D. and Parrinello, M. (2007) Canonical sampling through velocity rescaling. *The Journal of chemical physics*, **126**.
20. Berendsen, H.J., Postma, J.v., Van Gunsteren, W.F., DiNola, A. and Haak, J.R. (1984) Molecular dynamics with coupling to an external bath. *The Journal of chemical physics*, **81**, 3684-3690.
21. Parrinello, M. and Rahman, A. (1981) Polymorphic transitions in single crystals: A new molecular dynamics method. *Journal of Applied physics*, **52**, 7182-7190.
22. Barker, J.A. and Watts, R.O. (1973) Monte Carlo studies of the dielectric properties of water-like models. *Molecular Physics*, **26**, 789-792.
23. Sutthibutpong, T., Harris, S.A. and Noy, A. (2015) Comparison of molecular contours for measuring writhe in atomistic supercoiled DNA. *Journal of chemical theory and computation*, **11**, 2768-2775.
24. Song, Y., Kim, M., Sung, B.J. and Kim, J.S. (2025) Computational Characterization of DNA Catenanes. Journal of Chemical Theory and Computation, 21, 9967-9981.
25. Mitchell, J., Laughton, C. and Harris, S.A. (2011) Atomistic simulations reveal bubbles, kinks and wrinkles in supercoiled DNA. *Nucleic acids research*, **39**, 3928-3938.
26. Irobalieva, R.N., Fogg, J.M., Catanese Jr, D.J., Sutthibutpong, T., Chen, M., Barker, A.K., Ludtke, S.J., Harris, S.A., Schmid, M.F. and Chiu, W. (2015) Structural diversity of supercoiled DNA. *Nature communications*, **6**, 8440.
27. Pyne, A.L., Noy, A., Main, K.H., Velasco-Berrelleza, V., Piperakis, M.M., Mitchenall, L.A., Cugliandolo, F.M., Beton, J.G., Stevenson, C.E. and Hoogenboom, B.W. (2021) Base-pair resolution analysis of the effect of supercoiling on DNA flexibility and major groove recognition by triplex-forming oligonucleotides. *Nature Communications*, **12**, 1053.

28. Young, M.A., Jayaram, B. and Beveridge, D. (1997) Intrusion of counterions into the spine of hydration in the minor groove of B-DNA: fractional occupancy of electronegative pockets. *Journal of the American Chemical Society*, **119**, 59-69.
29. Feig, M. and Pettitt, B.M. (1999) Sodium and chlorine ions as part of the DNA solvation shell. *Biophysical Journal*, **77**, 1769-1781.
30. Manning, G.S. (1978) The molecular theory of polyelectrolyte solutions with applications to the electrostatic properties of polynucleotides. *Quarterly reviews of biophysics*, **11**, 179-246.
31. Manning, G.S. (1979) Counterion binding in polyelectrolyte theory. *Accounts of Chemical Research*, **12**, 443-449.
32. Naskar, S. and Maiti, P.K. (2021) Mechanical properties of DNA and DNA nanostructures: Comparison of atomistic, Martini and oxDNA models. *Journal of Materials Chemistry B*, **9**, 5102-5113.
33. Seol, Y. and Neuman, K.C. (2016) The dynamic interplay between DNA topoisomerases and DNA topology. *Biophysical reviews*, **8**, 101-111.
